# Brain-wide activity-identity mapping of neural networks associated with prosocial motivation in rats

**DOI:** 10.1101/2023.12.10.570980

**Authors:** Keren Ruzal, Estherina Trachtenberg, Ben Kantor, Hila Flumin, Adin Roemer, Andres Crespo, Johannes Kohl, Inbal Ben-Ami Bartal

## Abstract

A prosocial response to others in distress is an important driver of behavior across social species. To investigate the neurobiological mechanism leading to prosocial behavior, we use a helping behavior test wherein rats may release a trapped conspecific by opening a restrainer door. To ensure rats were not acting for social interaction, a separation divider prevented post-release contact (“**separated**” test). Despite the divider, most rats consistently opened the restrainer, demonstrating prosocial motivation. Brain-wide c-Fos analysis conducted via our opensource software "Brainways", revealed activity in empathy-related regions, including the anterior cingulate and insular cortices. Nucleus accumbens activity, previously recorded during helping, was not significant in the **“separated” test**. Chemogenetic manipulations of the accumbens did not prevent helping, suggesting that its activity reflects contact seeking. Mapping of oxytocin and dopamine receptors on active cells revealed region-specific recruitment of these subpopulations, depending on the social context. Network connectivity analysis highlights context-dependent functional subcircuits.

## Introduction

Prosocial behavior is a critical promoter of the survival and thriving of social groups. In particular, a prosocial response to others in distress has important implications for genetic continuity ^1–3^, and has evolved in altricial species both in the context of parental care as well as the broader social group. The powerful drive to act for the benefit of others has also evolved to be socially selective and heavily biased towards ingroup members across species 4^-11^. One of the proximate motivating factors for a prosocial response to distress in humans is empathic arousal, which involves sharing the affective state of others, coupled with a motivation to improve their well-being ^12,13^. Growing evidence points to similarities in the neurobiological mechanisms underlying empathy across altricial species ^14–17^. These shared mechanisms have likely evolved to drive a prosocial response to the needs of others, by engaging affective and motivational circuits ^18–21^.

The mobilization of an individual to help distressed conspecifics is a complex behavioral response which includes several components, like affective arousal, attentional shift, and motor preparation and execution ^13^. An integrative methodological approach is required to uncover the associated neurobiological processes that involve multiple cell types, neuromodulators, and interactions between dispersed neural networks. To this end, we employ a brain-wide approach that aims to outline the neural activity associated with prosocial motivation in a rat Helping Behavior Test (HBT). This strategy allows for an unbiased view of the neural patterns associated with specific behaviors rather than focusing on pre-selected regions of interest. Additionally, we investigate the identity of the active cells by mapping the receptor distributions of neuromodulators. Combining activity-tagging with cell identity, allows to uncover the role of different neuromodulators along the active network. We have developed for this purpose an opensource AI-based software, Brainways, that permits automated atlas registration and quantification of neural activity based on fluorescent markers (i.e., c-Fos) on coronal slices, and outputs the pattern of neural activity associated with a specific behavioral condition, based on partial least square tests (PLS) and network analyses ^22^. The Brainways pipeline for identifying neural connectomes was used here to investigate the prosocial motivation network. We hypothesized that others’ distress elicits activity in a dispersed neural network implicated with caregiving, including sensory, affective, and motivational circuits that are dependent on region-specific neuromodulation by oxytocin (OXT) and dopamine (DA) ^23^.

The rat HBT is ideally suited to explore this biological mechanism. In this simple test, a rat can help another rat escape a trap by opening a restrainer door ^24^. Rats are highly motivated to release cagemates without any previous training or external reward ^25^. It was previously shown that during the HBT, rats activate a neural network that is highly similar to that described in human empathy ^13^, including activation in the anterior cingulate and insular cortices as well as other prefrontal and limbic regions ^26^. A distinct subset of this network, including the nucleus accumbens (NAc) and other reward-processing regions, was only active in the presence of trapped ingroup members (rats of a familiar strain) but not for trapped outgroup members (rats of an unfamiliar strain). As rats selectively helped ingroup members, this suggests that reward-related activation is required for helping to occur. Yet, as the helping test allows for social contact following the release of the trapped rat, door-opening in the HBT could be explained by this social reward rather than being a prosocial act intended to help the conspecific.

To examine this possibility, a modified version of the HBT was established, where a divider separates the two rats after door-opening, thus dissociating the “social reward” experienced by social interaction from the “prosocial reward” experienced by helping. Therefore, a failure to release the trapped rat in the “separated” HBT would provide a strong indication that social reward is required for door-opening, whereas consistently releasing the trapped rat would point to a prosocial motivation to improve the other’s well-being. Moreover, neural activity of helpers in the “separated” HBT would reflect the prosocial response to the conspecific’s distress, dissociated from the expectation of social reward.

Thus, in this study, rats were tested in the “separated” HBT with trapped ingroup or outgroup members, and the neural network associated with these conditions was outlined. To further our understanding of the neural signature of prosocial motivation, the receptor distributions of OXT and DA were examined along with neural activity via multiplex RNAscope, as these systems hold particular importance for prosocial behavior ^27,28^. Considerable research, both in rodents and in humans, showed the involvement of the oxytocinergic system in empathy and prosociality ^29–35^, and in social bias ^36,37^. Similarly, activity in the mesocorticolimbic pathway has been associated with prosocial approach, pair-bonding, and altruism ^38–41^. Furthermore, OXT-DA dynamics specifically support pair-bonding and parental behavior in rodents ^42,43^ and thus likely influence prosocial motivation. More generally, OXT modulates dopaminergic pathways in the mesocorticolimbic system, hence affecting reward-related behaviors ^44^. Mapping the receptor distribution of the neurons active during the "separated" HBT would help identify whether parts of the prosocial motivation network were specifically sensitive to one or more of these neuromodulatory systems, and reveal regions of interaction. In addition, this analysis would indicate whether participation in the HBT shifted the receptor distribution compared to untested animals.

Here, converging evidence shows that helping behavior was preserved even in the absence of social reward. A distinct neural network associated with the “separated” HBT was identified via c­Fos immunofluorescence and RNAscope essays. The regions participating in the “separated” HBT comprised a subset of the original HBT network. The NAc, a central hub of the original HBT ^26^, was absent in the “separated” HBT network. To investigate this finding, we conducted a series of chemogenetic manipulations of the NAc, and found that helping was not reduced by NAc stimulation or inhibition. This suggests that NAc activity was related to the expectation of social reward rather than prosocial motivation. Discrete OXTr+ and DrD2+ subpopulations were involved in prosocial motivation and were modulated by the social identity of the conspecific in distress. In addition, network analysis based on the RNAscope dataset showed that each neuromodulatory system consists of distinct functional subpopulations, and highlighted potential pathways for further causal investigation.

## Results

### Rats consistently release a trapped rat even without post-release contact

Male Sprague Dawley (SD) rats were tested in the "separated” HBT (Fig. 1A), either with a trapped ingroup member (adult male cagemate, n = 8), or an outgroup member (adult male Long-Evans (LE) rat, n = 8). Over the 12 days of testing, most rats in the ingroup condition became “openers”, consistently opening the restrainer on the last 3 sessions and releasing the trapped rat (n = 5/8 openers, 62.5%, Fig. 1B). In line with this, the average door-opening latency significantly decreased for this group over days of testing (Friedman test, x^2^(11) = 21.33, p = 0.03, Fig. 1C). In the outgroup condition, a minority of rats became openers (n = 3/8 openers; 37.5%), and no significant change was recorded in the door-opening latency (p > 0.05). No significant difference was found between social conditions (p > 0.05 for these parameters). Although movement levels were similar across conditions (p > 0.05, Fig. 1D), rats in the ingroup condition spent more time near the trapped rat compared to the outgroup condition, (t(14) = 3.16 p = 0.01, Fig. 1E), indicating a trapped ingroup member was more salient. Baseline trait anxiety isn’t a likely alternative explanation of these results, as no difference was observed in open-field behavior between the groups (Fig. S1A-C). In sum, rats displayed door-opening behavior even in the lack of post-release social contact with the trapped rat, demonstrating that social reward is not necessary for helping to occur.

**Figure 1.**
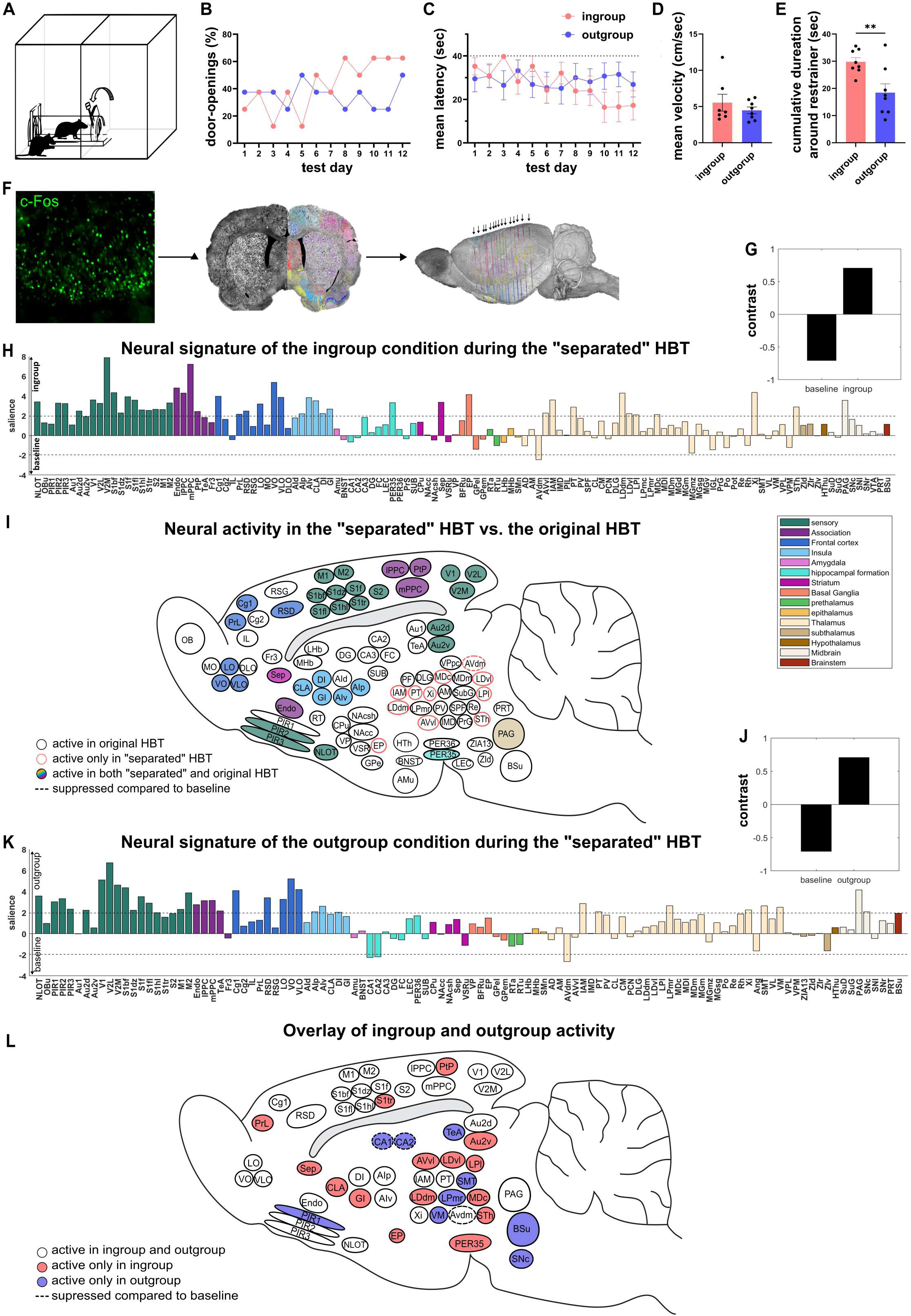
Neural signature of the “separated” HBT. **(A)** Diagram of the "separated" HBT paradigm showing the trapped rat released into a separate space. **(B)** The percentage of door­opening along testing days indicates most rats learned to open the restrainer. (C) Latency to door-opening reflects most rats opening the restrainer before the halfway point (dashed line). **(D-E)** Although activity levels were similar, rats in the ingroup condition spent significantly more time around the restrainer containing the trapped rat (mean ± SEM, **p<0.01). **(F)** Diagram of c-Fos registration and quantification via the Brainways software. **(G-H)** Neural pattern associated with the ingroup condition. Salience represents the z-score of bootstrapping tests. Regions crossing the threshold contributed significantly to the contrast (black bars). Arrows along the y-axis indicate the condition for which the region was more active. NAc activity did not contribute to the contrast. (I) Regions active during the "separated" HBT (in colors) are overlayed with the original HBT network (in white). **(J-K)** Neural pattern associated with the outgroup condition derived by PLS task analysis. **(L)** Overlay of regions active in the “separated” HBT ingroup and outgroup conditions. Identity-invariant regions are in white, and condition-specific regions in color.

### Neural activity during the “separated” HBT reveal prosocial motivation network dissociated from social reward

To index the neural activity associated with the "separated" HBT, the immediate early gene c-Fos was quantified following the final HBT session, where the restrainer was latched shut such that all rats experienced a full session with a trapped conspecific. Analysis of movements during the first 10 minutes of the session showed that rats in the ingroup condition spent significantly more time around the restrainer than did rats in the outgroup condition (t-test, t(13) = 3.05, p = 0.01, Fig. S1D-E). As a baseline for a non-specific signal, samples were collected from naive, age- and weight-matched SD male rats (n = 8). Tissue stained for c-Fos was registered and quantified via our in-house opensource software “Brainways” ^22^, PLS task analysis was used to outline the pattern of neural activity associated with behavior during the final session ^26,45,46^. PLS analysis of the ingroup and baseline conditions revealed a significant latent variable (LV, p = 0.01, Fig. 1G; See Fig S1F for c-Fos quantification per region). Next, permutation and bootstrapping tests were conducted to identify the contribution of each brain region to the contrast (Fig. 1H). A dispersed network of regions emerged across the brain, including specific subregions of the sensory (olfactory, auditory, visual, somatosensory) and motor cortex, associative cortices, frontal cortex (cingulate, orbital), insular, hippocampal, striatal, basal ganglia, thalamic, and midbrain regions. Notably, the identified regions in the “separated” HBT comprise a subset of the previously reported HBT network ^26^, which included a social reward component in the form of post-release contact (Fig. 1I).

Next, the pattern of activity associated with the outgroup condition was investigated. A significant LV emerged for the outgroup compared to baseline conditions (LV, p = 0.01, Fig. 1J-K). Bootstrapping analysis showed that many significant regions of the outgroup condition overlapped with the ingroup condition, including the anterior cingulate cortex (ACC), anterior insula (AI), sensory, and thalamic regions (Fig. 1L; see Table S1 for full list of regions). Thus, overall, a similar neural pattern is involved in the response to a trapped conspecific regardless of social identity. Yet some differences emerged, as the temporal association cortex (TeA), substantia nigra (SN), CA1 and CA2 of the hippocampus and layer 1 of the piriform cortex (PIR1) were selectively modulated in the outgroup condition (Fig. 1L). Conversely, the prelimbic cortex (PrL) and septum (Sep) were significantly modulated only in the ingroup condition (Fig. 1L). This is in line with previous findings in the original HBT, where the Sep, PrL, NAc and medial orbitofrontal cortex (OFC) were significantly more active for the ingroup condition compared to the outgroup condition ^26,47^. However, here, the PLS analysis shows that the NAc did not have a significant contribution to the ingroup condition. This was validated with an ANOVA (NAc-core, F(2,19) = 0.01, p = 0.99; NAc-shell, F(2,19) = 1.08, p = 0.36).

In sum, these data show that brain regions related to empathy and social cognition were active in the “separated” HBT. The lack of NAc activity may reflect the lack of expectation for social reward. The experiments below further validate and expand on this unexpected finding.

### OXT and DA receptor distribution mapping on active cells

Next, we set out to map oxytocin (OXT) and dopamine (DA) receptor distributions within c-Fos^+^ cells. Adult LE rats were tested with trapped ingroup (n = 16 males, n = 16 females) and outgroup (n = 15 males, n = 8 females) members in the “separated” HBT (Fig. 2A). In the ingroup condition, 53% of the rats became openers (n = 10/16 male openers, n = 7/16 female openers). Similarly, 61% of rats released trapped outgroup members (n = 8/15 male openers, n = 6/8 female openers, Fig. 2B). Opening latencies were significantly reduced in all four groups along testing days, indicating an overall increase in helping behavior as more rats learned to open the restrainer (Friedman’s, p < 0.05; Fig. 2C). Yet, on the last 3 days of testing, when opening patterns stabilized, a significant difference emerged between the mean % door-opening (t-test, t(1,10) = 4.65, p < 0.01) and opening latency (t-test, t(1,10) = 3.98, p < 0.01), showing a mild effect of social condition on prosocial behavior. Analysis of movement levels showed no main effect for sex or social condition (ANOVA, p > 0.05, Fig. 2D), yet a significant interaction emerged (F(1,51) = 4.47, p = 0.04) as females tended to be more active for ingroup members whereas males tended to be more active in the outgroup condition. Moreover, males spent significantly more time around the restrainer containing the trapped ingroup member than the three other groups (ANOVA, p < 0.05; Bonferroni, p < 0.01; Fig. 2E). Therefore, ingroup bias may be more prominent in males, who also tended to open less for an outgroup member. This study replicates the finding above showing that most rats are motivated to release trapped conspecifics even in the absence of access to the released rat.

**Figure 2.**
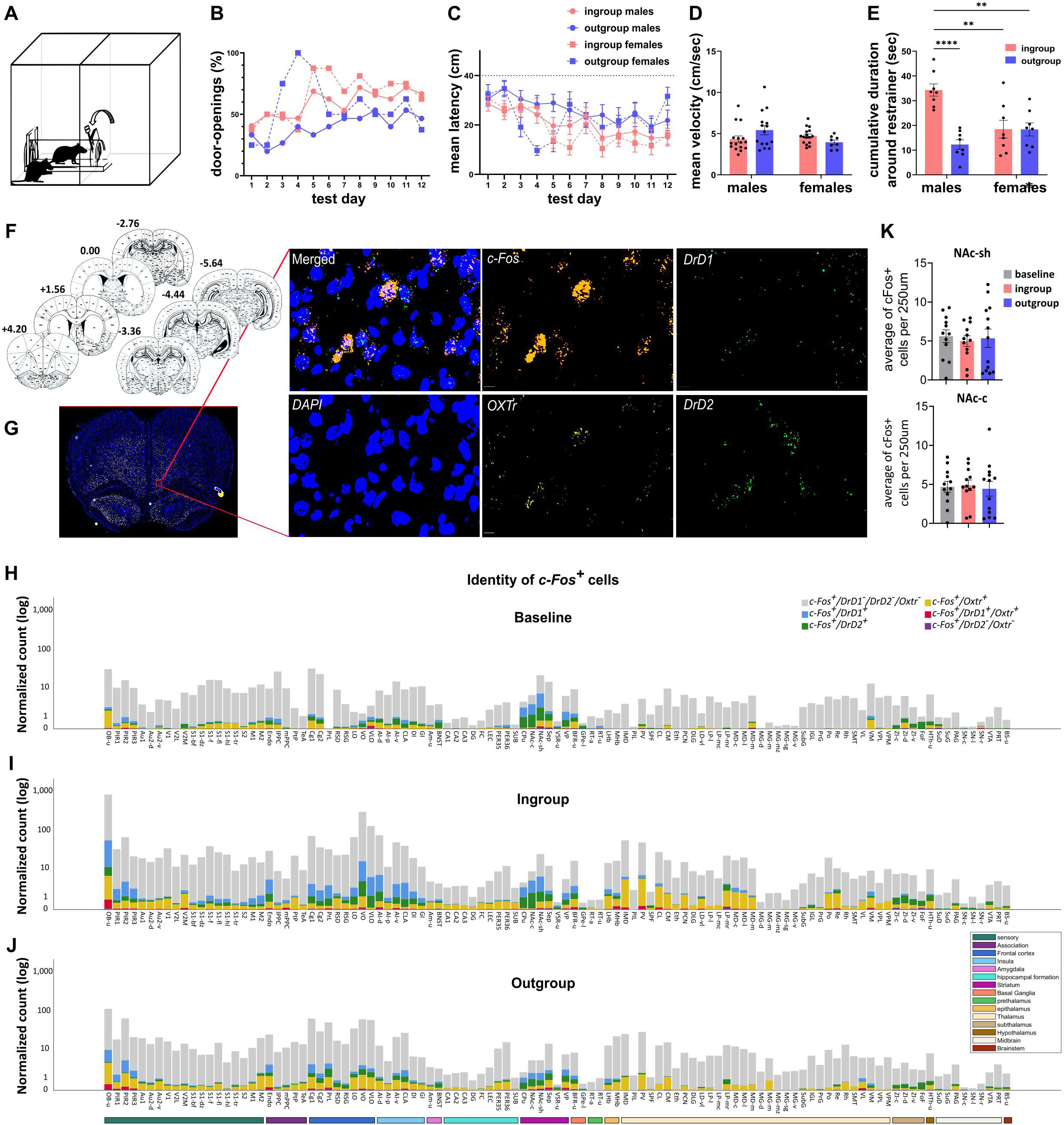
Receptor mapping on active cells during the "separated" HBT. **(A)** Diagram of the behavioral setup **(B-C)** Door-opening percents and latencies show about half of the rats became openers in the ingroup and outgroup conditions. Dashed line represents the halfway door-opening point. Movement analysis of velocity **(D)** and time around the restrainer **(E)** show that males and females treat outgroup members differently (means ± SEM). **(F-G)** Coordinates and sample images of the histological images used for quantification of c-Fos, OXTr, DrDl, and DrD2. **(H-J)** Distributions of DA and OXT receptors within the c-Fos+ population in the baseline, ingroup, and outgroup HBT conditions, respectively. (K) quantification of c-Fos in the NAc across experiments shows no effect of condition. **p<0.01; ****p<0.0001.

To identify the receptor distribution along the active (c-Fos^+^) network, quantification of mRNA for c-Fos, DrD1, DrD2, and OXTr was conducted via RNAscope in a subset of these rats (n = 6 ingroup; n = 6 outgroup; 3 males and 3 females per condition; Fig. 2F-G). A baseline condition of untested rats was used to contrast out non-specific signal ("baseline"; n = 5). Receptor distribution within *c-Fos^+^* cells varied across brain regions, in a condition-dependent manner (Fig. 2H-J). *c-Fos^+^/DrD1^+^* and *c-Fos^+^/DrD2^+^* subpopulations were especially apparent in the striatum across conditions, comprising on average 12.44 ± 9.53 and 21.18 ± 9.82 percent of total *c-Fos^+^* cells respectively. Additionally, *c-Fos^+^/OXTr^+^* was prominent in the amygdala (6.39 ± 5.89) hippocampus (4.97 ± 5.82), insula (4.06 ± 4.64) and striatum (4.57 ± 3.23) (See Table S2 for all regions). Typically, cells were sensitive to no more than a single receptor, with only 0%-5% of *c-Fos^+^* cells co-labeled for two or more receptors, indicating that most active cells were recruited from distinct populations. PLS analysis of *c-Fos* for the ingroup and baseline conditions revealed a significant LV (p = 0.046, Fig. S2). Similarly to the first study, NAc activation did not differ from baseline levels (Fig. 2K), further evidence for the lack of NAc involvement in the “separated” HBT.

To find the subcircuit of active cells that are sensitive to OXT or DA signaling, we looked at co­labeled cells for c-Fos and each receptor. PLS analysis did not find significant LVs in the normalized numbers of any of the subpopulations compared to baseline. Next, to examine to what extent each subpopulation was recruited during the task, PLS task analysis was conducted on the percent of each co-labeled subpopulation. A significant LV would indicate that the ratio of these subpopulations was affected by the social condition.

### DA modulation during the “separated” HBT is driven by DrD2+ cells on the active population

PLS analysis of the percent of *c-Fos^+^/DrD1^+^* cells did not indicate a brain-wide difference from baseline levels (PLS LV, p > 0.05). However, the percent of *c-Fos^+^/DrD2^+^* cells in the ingroup condition did show a global difference from baseline (PLS LV, p = 0.016; Fig. 3A). This contrast was driven by a dispersed set of regions, such as secondary auditory and visual regions (Au2d, V2L), PrL, retrosplenial (RSG), TeA, insular subregions (AId, GI), the ventral pallidum (VP) and the deeper layers of the superior colliculus (SuD). The Fields of Forel (FoF) and Superficial gray layer of the superior colliculus (SuG) were suppressed compared to the baseline condition (Fig. 3A). PLS of the outgroup and baseline conditions for the %c-Fos+/DrD2+ subpopulation also revealed a significant LV (p = 0.048), with a significant contribution of the Au2d, RSG, VP, ventral striatal region (VSR), lateral habenula (LHb), periaqueductal gray (PAG). Several regions were significantly suppressed compared to baseline including the PIR3, endopiriform nucleus (Endo), posterior AI (AIp), and FoF (Fig. 3B).

**Figure 3.**
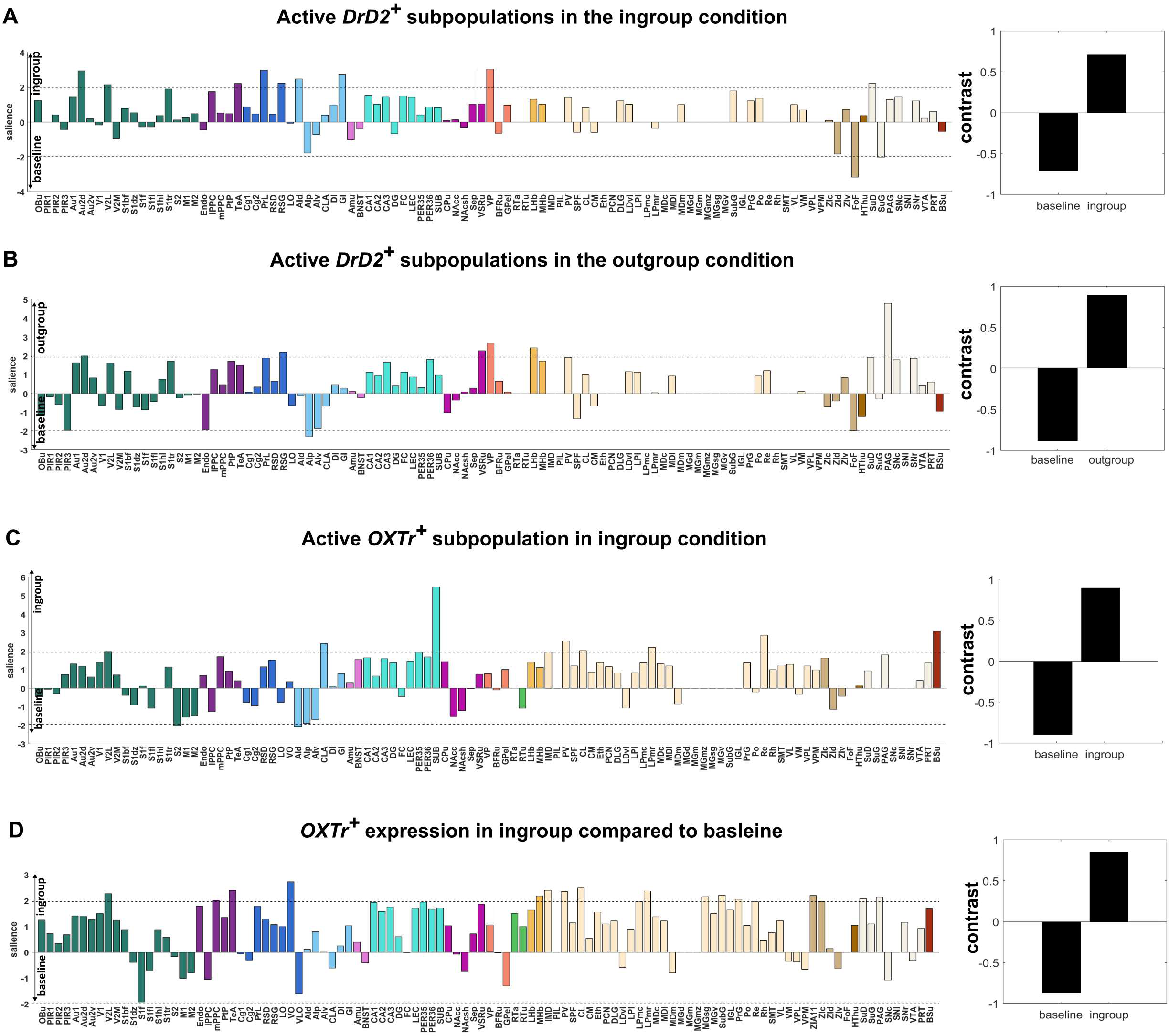
PLS analysis of receptor-specific subpopulations of active cells in the "separated" HBT. A significant LV indicates a brain-wide effect of condition on %c- Fos+/DrD2+ subpopulations for the ingroup **(A)** and outgroup **(B)** conditions. Black line denotes significance threshold. **(C-D)** OXT showed significant modulation for the ingroup condition, with a significant LV for the %c-Fos+/OXTr+ subpopulation as well as number of OXTr+ cells (independent of c-Fos) in the ingroup compared to baseline. All regions that cross the black threshold line contributed significantly to the contrast.

### Participation in the “Separated” HBT causes a shift in OXTr expression for the ingroup condition

For the ingroup, PLS analysis found a significant LV for *%c-Fos^+^/OXTr^+^* (PLS LV, p = 0.046; Fig. 3C), indicative of a brain-wide pattern associated specifically with this condition. The V2L, claustrum (CLA), SUB, brainstem and thalamic subregions (IMD, PV, LPmr, and Re), S2 and dorsal AI (AId) contributed significantly to this contrast. The S2 and AId were suppressed in the ingroup compared to baseline levels. Interestingly, AI suppression was observed for the *c-Fos^+^* DrD2 as well as OXTr for the ingroup condition.

Independently of c-Fos, a brain-wide increase was observed for *OXTr^+^* cell numbers in the ingroup condition compared to baseline (LV, p = 0.049, Fig. 3D). Significant upregulation of OXTr was observed in the V2L, mPPC, TeA, VO, MHb, and several thalamic, midbrain and brainstem subregions (Fig. 3D). This suggests participation in the “separated” HBT ingroup condition for two weeks modulated OXT receptor expression in various brain regions, including in cells that were not active during the task on the c-Fos quantification day. A similar analysis of the *DrD1+* and *DrD2+* cell numbers was not significant, yet focal changes are not captured by this global analysis.

Finally, in some regions both OXT and DA were sensitive to the social condition. An intersection was observed in the AI, V2L, TeA, SuD, PV, and PAG (Fig. 4A). These areas are likely targets for hubs that merge information from these two systems. In sum, these combined data demonstrate that neural activity related to the “Separated” HBT relies on largely separate systems for OXT and DA signaling, with specific regions where modulation includes both systems and point to the centrality of the AI for these neuromodulators.

**Figure 4.**
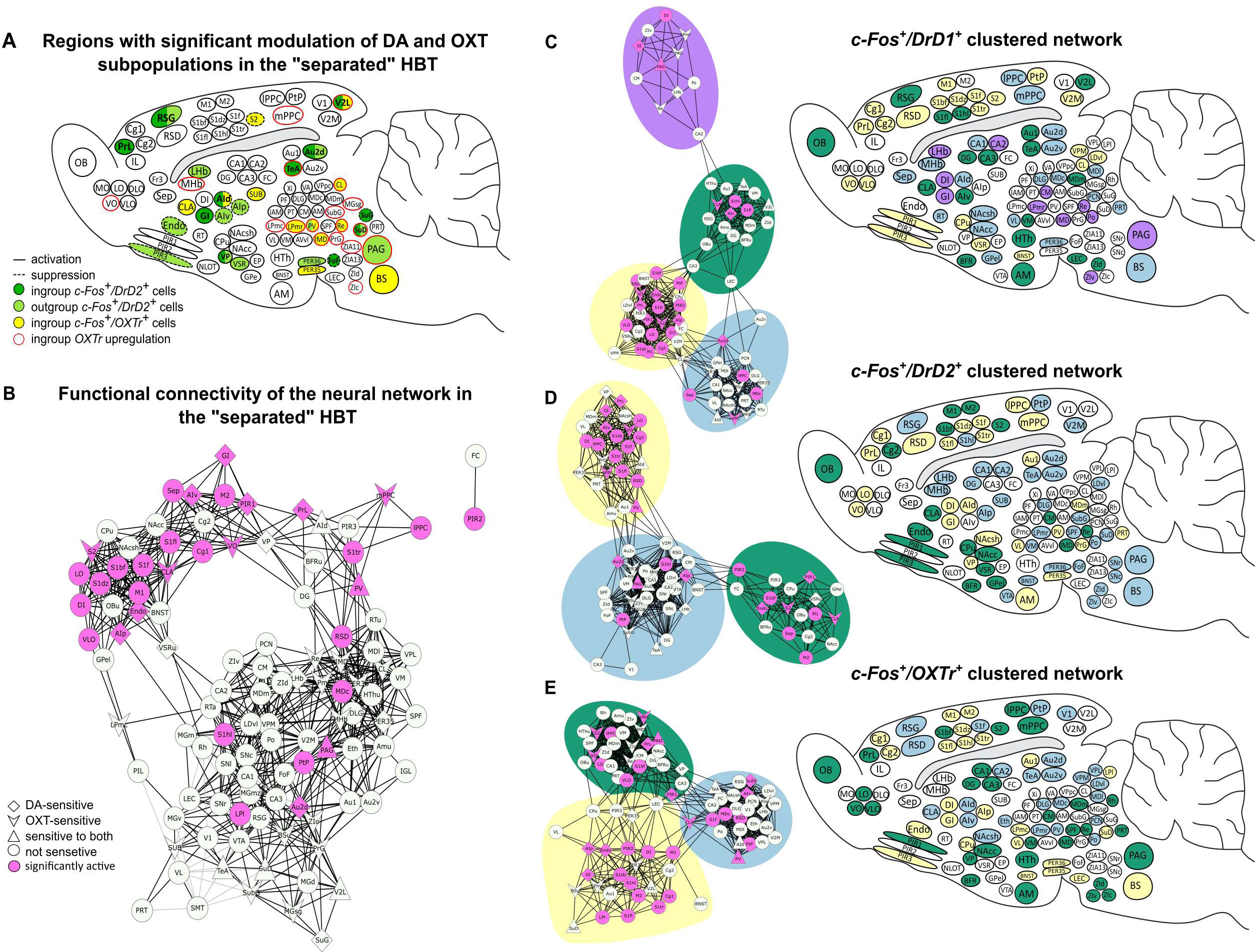
Neural networks underlying prosocial motivation. **(A)** Summary diagram of brain regions showing significant modulation in DA and OXT receptor subpopulations, based on PLS analyses presented in Fig. 3. Most brain regions were sensitive to either DrD2+ or OXTr+, whereas a few regions, including the AI, were sensitive to both. (B) Network graph based on c- Fos+ cell numbers depicts the functional connectivity for the “separated” HBT ingroup condition. Lines between brain regions (edges) represent significant Pearson’s correlations (p < 0.05, positive correlations in black, negative in gray). A main cluster contains most of the significantly active regions (pink). **(C-E)** On right, network graphs of the active subpopulations within each neuromodulatory system. The active network (pink) is displayed, with receptor sensitivity denoted by shape. Background color represents clusters identified via a Louvain algorithm. On right, brain diagrams summarizing functional clusters shown.

### Distinct functional OXT and DA clusters in the active network associated with the “separated” HBT

To gain insight into the brain-wide functional connectivity during the HBT, network graph theory was used to generate maps based on *c-Fos* quantification from each social condition in the “separated” HBT. The network graphs were generated from inter-region Pearson’s correlation matrices, thresholded at p < 0.05 and clustered using a Louvain algorithm for community detection (see methods, Fig. S3A-C). In the ingroup condition, a network graph based on *c-Fos^+^* cell numbers was comprised of five clusters (Fig, S3D). One central cluster was composed mostly of regions that were significantly active during the “separated” HBT (main cluster: 59% of active regions, less than 16% for each of the other clusters; Fig. 4B, Fig, S3D). The strong connectivity of these regions provides further indication for their role in prosocial motivation. Notably, although significant activity was not found in the NAc (see Fig. 2K), it was highly connected to other regions in this main cluster. This pattern was not maintained for the outgroup or baseline networks (Fig. S3E-F).

Next, to investigate the connectivity within each neuromodulatory system, network graphs were generated based on cell numbers of co-labeled subpopulations for *c-Fos and DrD1, DrD2*, or *OXTr* (Fig. 4C-E). For the ingroup condition, the subnetworks were highly clustered and dense, suggestive of functional sub-circuits (Fig. 4C-E; Clustering co-efficient: c-Fos^+^: 0.65; *c-Fos^+^/DrD1^+^*: 0.80; *c-Fos^+^/DrD2^+^*: 0.79; *c-Fos^+^/OXTr^+^*: 0.78. Density: *c-Fos^+^*: 0.14; *c-Fos^+^/DrD1^+^*: 0.21; *c-Fos^+^/DrD1^+^*: 0.23; *c-Fos^+^/OXTr^+^*: 0.23). For DA, one main cluster contained many of the regions significantly active during the "Separated" HBT ingroup condition compared to baseline *(c-Fos*^+^*/DrD1*^+^: 53%, 16%, 13% per cluster; *c-Fos*^+^*/DrD2*^+^: 47%, 28%, 16%; Fig. 3C-D). In the oxytocinergic subpopulation, the “separated” HBT network was dispersed across three clusters (41%, 31%; 25% Fig. 4E).

It is possible that the unique modulatory structure of the receptor subnetworks reflects sampling of the same parent network. However, each network had unique connectivity, and the main cluster included different active regions. Alternatively, the modularity could result from condition­independent characteristics related to the receptor distribution. Yet, in networks generated based on all rats across conditions (“all” network), modularity was significantly lower (mean modularity for “all”: 0.32 ± 0.02, ingroup: 0.59 ± 0.02, t-test, p < 0.01), and inter-region correlations were weaker overall (Fig. S3G-L). This suggests that the highly segregated dense clusters observed in the ingroup condition are context-dependent. Future experiments manipulating these sub-circuits are needed to uncover their distinct functionalities. We conclude that the oxytocinergic and dopaminergic systems should not be considered as holding a unified function in the brain.

### Door-opening is not altered by NAc modulation

As demonstrated above, the role of the NAc in helping behavior is not clear. In the two studies above, activation was not observed in the NAc for the ingroup condition in the “separated” HBT beyond levels seen in the baseline or outgroup conditions (Fig. 2K). Yet it was an integral part of the functional network (Fig. 4B). To investigate directly whether NAc activation was required for helping to occur, NAc activity was manipulated chemogenetically in adult male LE rats tested with a trapped ingroup member (an LE stranger) in the original, non-separated HBT (Fig. 5A). Bilateral injections of designer receptors activated only by designer drugs (DREADDs) to the NAc (A/P: 1.5, M/L: ±1.5, D/V: -8.0) were performed three weeks prior to testing of either an inhibitory (n = 8, AAV8-hSyn-hM4D(Gi)-mCherry; “inhibition” condition), or excitatory (n = 8, AAV8-hSyn-hM3D(Gq)-mCherry; “excitation” condition) DREADD (Fig. 5B; summary of viral spread Fig. S4A-B). The ratio of c-Fos+ neurons was successfully modulated by Clozapine-N-Oxide (CNO) administration (Fig. 5C-D), and viral infection was verified in all rats (Fig. S4A-B).

**Figure 5.**
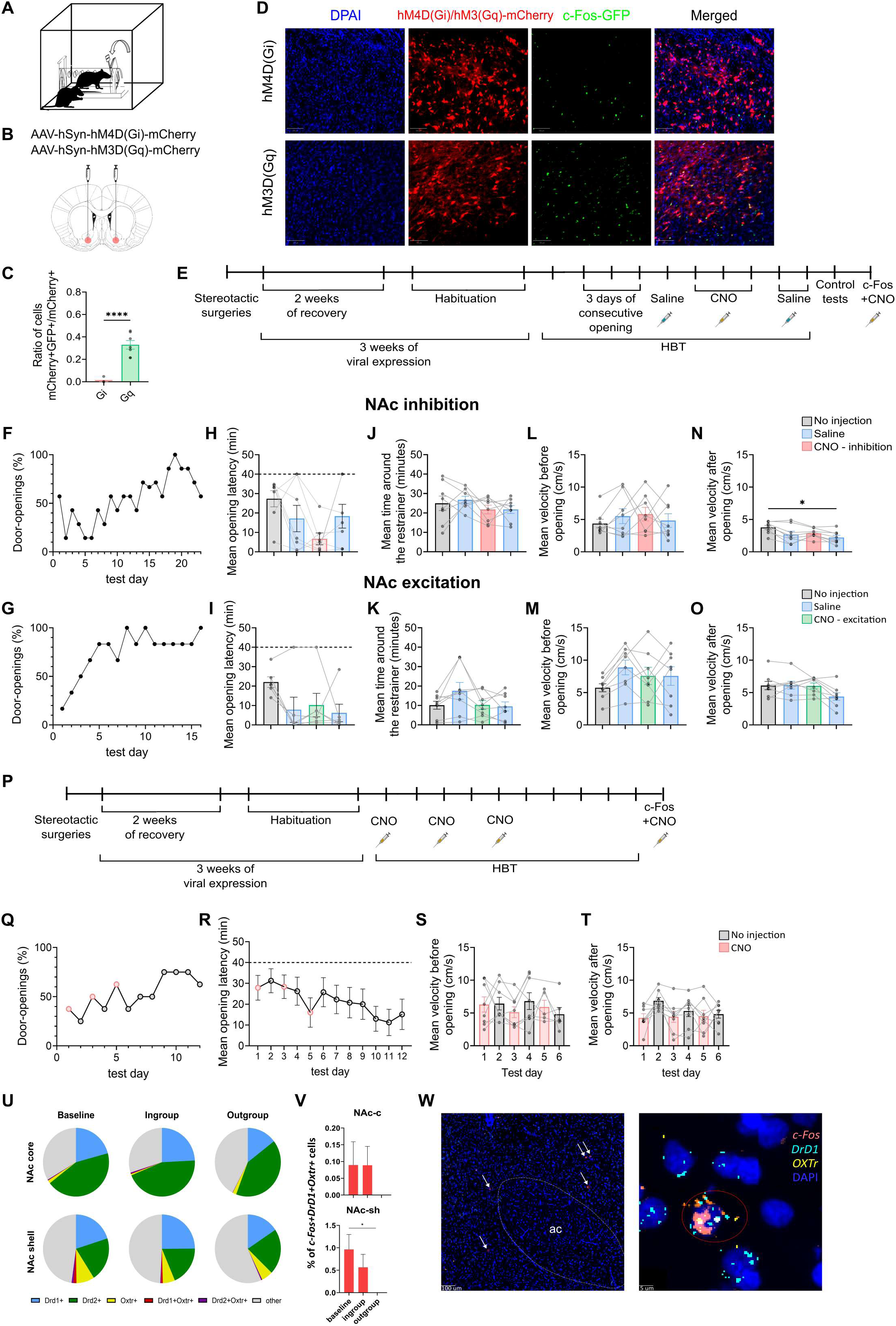
NAc activity is not necessary for maintenance or learning of door-opening behavior. **(A)** Rats were tested in the original (non-separated) HBT. **(B)** Neural activity in the NAc was inhibited or excited with DREADDs. **(C-D)** Co-labeling of the viral-infected cells with c-Fos as a validation of the chemogenetic manipulation via CNO administration. **(E)** Timeline of experiments 1-2: chemogenetic NAc manipulation in “expert” door-openers (after 3 consecutive openings). (F-O) behavioral results from the inhibition (top row) and excitation (bottom row) experiments indicate no effect of the manipulations on door-opening. Percentages of door-opening increased (F-G), and average door-opening latencies before the halfway point (dashed line) decreased **(H-l).** CNO did not alter time around the restrainer **(J-K)** or velocity before **(L-M)** or after **(N-O)** door-opening. **(P)** Timeline of experiment 3: chemogenetic manipulation in “naive” rats (from day 1 of the HBT). **(Q-R)** Manipulation did not prevent learning of door-opening as expressed by the percentages and latencies of door-openings. **(S-T)** No effect was recorded on velocity before or after door-opening. **(U-V)** receptor identity of c-Fos+ cells in the NAc. (W) Representative image of NAc co-expression of *c-Fos, DrDl,* and *OXTr.* “ac” = anterior commissure. Error bars represent SEM. *p<0.05, ****p<0.0001.

First, we examined whether NAc manipulation was required for maintaining door-opening in established openers. A rat was considered an “opener” after three consecutive door -opening days. The range of days to reach the criterion varied considerably between individual rats (1-15 days). One rat in the inhibition condition and two rats in the excitation condition never opened the restrainer and were excluded from the analysis. Once reaching the criterion, rats were treated with saline and then CNO (3 mg/kg), over the next five sessions (Figure 5E). Saline trials served to exclude behavioral changes due to the i.p. injection, which is a mildly stressful manipulation. We found that CNO administration did not impede door-opening in either condition as expressed by door-opening ratio and latency before, during, and after CNO administration (repeated ANOVA; p > 0.05; Fig. 5F-I). Similarly, no differences were observed in the time spent around the restrainer or activity levels before door-opening (ANOVA, p > 0.05; Fig. 5J-M). A significant decrease in activity levels was observed after door-opening in the inhibition condition (ANOVA, p < 0.05) but this effect was not driven by CNO administration, as no difference was observed between CNO and saline sessions (Fig. 5N). No effect on activity was observed for the excitation condition after door-opening (Fig. 5O). Finally, CNO did not affect motor activity, anxiety, or non­social reward-seeking or consumption in tests performed after the HBT on these same animals (Fig. S5).

Next, we sought to determine whether NAc activity was required for the initial acquisition of door­opening behavior. To this end, another cohort of adult male LE rats (n = 8) was tested in the original non-separated HBT under chemogenetic inhibition of NAc activity (AAV8-hSyn-hM4D(Gi)-mCherry, see Fig. S4C for viral spread). HBT testing began three weeks after virus injection, and CNO was administered in alternating sessions (days 1,3,5), and on the final session for histological validation (Fig. 5P). Most rats (n = 7/8) became openers (Fig. 5Q), and door-opening latency was significantly reduced along testing sessions (x^2^(11) = 28.382, p = 0.003, Fig. 5R). Analysis of activity levels shows did not find an effect for CNO either before or after door-opening on the first week of testing (Fig. 5S-T). In sum, these experiments show that general excitation or inhibition of the NAc does not impede door-opening acquisition or maintenance, in either inexperienced or established openers.

Finally, receptor mapping of *c-Fos^+^* cells in the NAc revealed that significantly fewer cells co­expressed *DrD1* and *OXTr* in the outgroup condition compared to the ingroup and baseline (F(1,7) = 9.63, p = 0.017, Fig. 5U-W). This finding was supported by PLS analysis (LV, p = 0.048, Fig. S6). Suppression of this subpopulation in the outgroup may be related to threat arousal in this social context and present an interesting target for future manipulations.

Collectively, evidence from the experiments presented above indicates that the NAc, while playing a role in the HBT, was not specific to prosocial motivation. While similar activity was found across social conditions, and NAc inhibition or excitation was not sufficient to prevent helping behavior, the NAc nonetheless was a central part of the functional network of the ingroup condition, and differences were observed in the active receptor-sensitive subpopulations according with condition.

## Discussion

This study explores the neural correlates of prosocial motivation dissociated from anticipation of social reward, and the neuromodulatory systems involved. We employed “activity-identity” mapping via c-Fos and receptor quantification with Brainways, a novel open-source AI-based software developed in-house. In order to examine prosocial behavior without an expectation for post-release contact, a “separated” HBT paradigm was used, where after door-opening the trapped rat escapes into a separate space. In contrast with a previous version of the "separated" HBT ^26^, here rats did not have any previous experience releasing a trapped rat and had not ever experienced the social reward associated with post-release contact. We found that the percent of rats who became helpers in the "separated" HBT was somewhat below that observed for the original HBT (60% vs. 70% openers). This is an indication that social contact, which is undoubtedly rewarding in some conditions, comprises only part of the motivation for door-opening in the original paradigm. As many rats consistently released the trapped conspecific, across strains, sexes, and group memberships even in the lack of social contact, it is not likely the primary driving factor for door-opening. This finding adds support to research demonstrating social contact is not required for helping ^35,48,49^, and promotes the idea that rats are motivated by a prosocial intention to improve the trapped rats’ wellbeing. The "separated" paradigm presented here is thus an improved behavioral test for targeted helping motivated by empathy to distress.

We found that in the "separated" HBT helping occurred, to a non-negligible extent, also towards trapped outgroup members. In previous experiments, helping was only observed towards outgroups in adolescent rats ^47^. This finding suggests that reducing post-release interaction can increase prosocial behavior towards outgroup members in rats. The underlying cause of this effect remains unclear. Eliminating contact with the released rat may have reduced threat arousal associated with post-release interaction, or reduced distress by increasing the distance between the two rats. As helping ingroup members was previously found to reduce corticosterone arousal ^50^, these motivations are difficult to disentangle, and will need to be examined in future studies. Alternatively, the effect could have been driven by strain and sex differences, where social bias is less pronounced in females.

The NAc was previously identified as a central network hub in the original HBT, and was shown to be active when rats approach a trapped ingroup member ^26,47^, leading to the interpretation that the NAc plays a role in prosocial motivation. Yet, analysis of neural activity in the "separated" HBT found no increase in NAc activity across two separate cohorts. Furthermore, helping was not prevented by NAc manipulation via chemogenetic excitation or inhibition. This combined evidence suggests that NAc activity previously observed during the HBT may represent social reward, as previously reported ^51^. However, it is important to note that even in the "separated" HBT, a sparse population of cells was active in the NAc, and it was part of the functional network of the ingroup condition. Thus, the role of this region requires further investigation.

The findings from the studies presented in this paper are added to previous published data, all showing converging evidence for similar regions activated during the HBT. These regions form a dispersed network across the brain, parts of which are activated only in a specific social context, and others which are recruited regardless of the trapped rats’ identity. The socially selective regions may play a role in social identity representation, and the ensuing cascade of affective and motivational implications. Thus, ingroup-specific activity is associated with prosocial motivation, whereas outgroup specific activity may reflect inhibition or threat arousal. The identity-invariant activity may represent processing of observed distress, which is expressed by trapped rats across conditions.

Analysis of the receptor distribution within the ensembles activated during the ingroup condition revealed elevated activity of *DrD2^+^* and *OXTr^+^* cells in a variety of regions, and a general upregulation OXTr compared to untested animals. The AI, of particular importance in the empathy network, was differentially modulated for ingroup and outgroup members in its different subregions, suggesting that although as a whole the AI was active in both social conditions, different subpopulations were involved. The oxytocinergic and dopaminergic systems were found to have another intersection - in the NAc. Rats undergoing the HBT in the outgroup condition recruited fewer cells co-expressing *OXTr* and *DrD1.* As rats don’t tend to have affiliative interactions with outgroup members ^47^, this finding is in line with literature documenting the involvement of *OXTr* and *DrD1* in the neurobiological mechanism of social interactions and approach ^52,53^. The exact mechanism underlying the contribution of these regions and neuromodulatory systems to prosocial motivation and behavior requires further investigation.

To expand our understanding of the neural patterns found above, functional connectivity networks were constructed for each social condition based on *c-Fos^+^* expression. Most of the regions comprising the “separated” HBT network were also functionally connected in the ingroup condition, and these connectivity patterns were altered in other social contexts. As many of the functional connections are in line with structural connectivity ^54^, these findings provide an abundance of targets for causal manipulations, to help determine the mechanism underlying prosocial motivation and behavior. For example, the OFC and PrL were functionally connected only in the ingroup condition. Moreover, while this connection remained for all three sub-networks, PLS analysis suggests the *DrD2^+^* is especially important. The AId-NAcsh is another interesting projection, in which connectivity was robust across receptors and social contexts, and which is modulated by both DA and OXT. This projection may encode the presence of conspecific distress. Furthermore, ACC-NAc *DrD2* connectivity was specific to the ingroup condition, supporting previous reports of its role in empathy in rodents ^16’26^. It’s important to keep in mind that other neuromodulators likely participate in prosocial behavior. For example, sex hormones like testosterone or estradiol are involved in parental behaviors ^55^ and should be examined in future studies in a broad prosocial context.

The c-Fos data is based in one dataset on immunofluorescence and in another dataset on RNAscope, inserting additional variability to the neural analysis. c-Fos mRNA levels are maximally expressed 30 minutes after the salient event’s onset 56, whereas the immunofluorescence data reflects activity taking place in the previous hour. Thus, some differences in these datasets could be explained by the underlying methodology. As many commonalities emerged, and as the emphasis in the RNAscope data was on receptor mapping, combining these methods appears to have enhanced the results. Another limitation of this study is the small sample size in the RNAscope analysis - each social condition comprises six rats. As males and females were combined, sex differences in the neural circuits encoding prosocial motivation may not be revealed. Finally, the reliance on c-Fos as a marker for neural activity has several caveats, as we previously described ^26^. Besides having low temporal resolution, Fos is an indirect indicator of activity and may represent information about plasticity or synchrony along with spiking. Furthermore, the reliance on c-Fos as a neural activity marker allows for only correlational results. Thus, the network presented here should be further investigated using causal manipulations of neural activity.

> In conclusion, this study outlines the neural network associated with the motivation to help others in distress in rats, dissociated from activation due to social reward. The complex interconnections of the OXT and DA systems during prosocial motivation are highlighted and shown to underlie distinct networks of neural functional clusters. The unbiased brain-wide “activity-identity” mapping approach we employ here is highly informative and reveals new research directions for the elucidation of the neural mechanism underlying empathic helping behavior and ingroup bias.

## Acknowledgements

We thank Dr. Einat Bigelman for assisting with the RNAscope collaboration, and Prof. Cornelius Gross for valuable input. We are also grateful to Tamar Spectre, Nesim Gnceer, Aviva Polisar, Annaelle Bismuth, Nir Doron, Or Peleg, Shir Toledano, and Tal Bar-Nahor for their help with data collection. This work was funded by the Israel Science Foundation and the Azrieli Foundation.

## Author Contributions

Conceptualization, I.B.B. and K.R.; Investigation, K.R., E.T., H.F., A.R.; Analysis, I.B.B., K.R., E.T., B.K.; Visualization & Writing, I.B.B. and K.R.; Methodology, I.B.B., J.K., A.C; Resources & Funding, I.B.B; Supervision, I.B.B. and J.K.

## STAR★Methods

### Key resources table

| REAGENT or RESOURCE | SOURCE | IDENTIFIER |
| --- | --- | --- |
| **Antibodies** |  |  |
| Rabbit anti-c-Fos primary antibody | Abcam | Abcam Cat# ab190289, RRID:AB_2737414 |
| Donkey anti-Rabbit IgG (H+L) Highly Cross-Adsorbed Secondary Antibody, Alexa Fluor™ 488 | Thermo Fisher Scientific | Thermo Fisher Scientific Cat# A-21206, RRID:AB_2535792 |
| **Deposited data** |  |  |
| https://osf.io/ve8s7/?view_only=e8880e1eb934416db17ad5eaf8a9d7da | N/A | This paper |
| **Experimental models: Organisms/strains** |  |  |
| Sprague-Dawley Rat | In-house breeding | N/A |
| Long-Evans Rat | In-house breeding | N/A |
| **Software and algorithms** |  |  |
| MATLAB | Mathworks (https://www.mathworks.com/) | RRID: SCR_001622 |
| Python | Python Programming Language http://www.python.org/ | RRID:SCR_008394 |
| SPSS | IBM | RRID:SCR_019096 |
| GraphPad Prism | http://www.graphpad.com/ | RRID: SCR_002798 |
| QuPath | https://qupath.github.io/ | RRID:SCR_018257 |
| Cytoscape | https://cytoscape.org/ | RRID:SCR_003032 |
| Graph-tool | https://graph-tool.skewed.de/ | N/A |
| **Plasmids** |  |  |
| pAAV5-hSyn-hM4D(Gi)-mCherry | http://www.addgene.org/50475 | RRID:Addgene_50475 |
| pAAV5-hSyn-hM3D(Gq)-mCherry | http://www.addgene.org/50474 | RRID:Addgene_50474 |
| **Chemical compounds** |  |  |
| Clozapine N-oxide | Tocris Bioscience | Cat. No. 4936 |
| **Other** |  |  |
| FISH probe: Fos | ACDBio | Cat No. 403598 |
| FISH probe: OXTr | ACDBio | Cat No. 483678 |
| FISH probe: DrD2 | ACDBio | Cat No. 315648 |
| FISH probe: DrD1a | ACDBio | Cat No. 317038 |

### Resource availability

#### Lead contact

Further information and requests for resources and reagents should be directed to and will be fulfilled by the lead contact, Inbal Ben-Ami Bartal.

#### Materials availability

This study did not generate any new reagents or animal lines.

### Experimental model and subject details

#### Animals

Rat studies were performed in accordance with protocols approved by the Institutional Animal Ethics Committee at Tel-Aviv University, Israel. Rats were socially housed in cages of two same­sex individuals, in a temperature (22-24C) and humidity controlled (55% relative humidity) animal facility, on a 12:12 light:dark cycle (lights on at 07:00). Food and water were provided ad libitum. All testing was done in the rat’s light cycle. In total, 16 male Sprague-Dawley rats, 56 male Long Evans rats, and 24 female Long Evans rats were tested as free rats across all experiments; brains from these animals were collected for subsequent processing. The same amount of rats was used as trapped rats across all experiments. All rats were bred in our animal facility. Animals were separated by sex and weaned at p21, then were housed in pairs one week later at p28. Trapped and free rats were of the same sex and age. All rats tested were 2-8 months old. Sprague Dawley/Long Evans animals were assigned to one of two experimental groups: they were either tested with cagemates (a rat from the same strain; ingroup) or with strangers (a rat from the other strain; outgroup). In the outgroup condition, the trapped rats, as well as the free rat, had never been exposed to a rat of the other strain before the first day of testing. The only exposure was such that was afforded by the 1h testing sessions for the two weeks of behavioral testing, which was previously shown to be insufficient to lead to door-opening ^25^. All rats were pair-housed. Rats in the ingroup conditions were housed with the trapped cagemate they were encountering during the sessions of the helping behavior test. Rats in the outgroup condition were housed with a cagemate who was also tested as a free rat in the helping behavior test with a stranger of the other strain. In the Chemogenetic experiments, Long Evans rats were tested with an unfamiliar ingroup member (i.e., not a cagemate).

### Method details

#### Apparatus

All behavioral procedures were conducted in 52X52X52 plexiglass chambers under light conditions of 80-120 lux. In the new "separated” HBT presented here, a plexiglass wall (transparent and perforated at the bottom) was used to divide the arena into two equal compartments. All behavioral apparatuses were cleaned with 1% acetic acid and general cleaning soap after every behavioral procedure.

#### Behavioral Testing

##### Habituation

Animals underwent five days of habituation, which included daily boldness test sessions followed by 5-minute handling sessions and 30-minute habituation sessions in the arena. On the first day, rats were first tested in an Open Field test (OF) for 15 minutes and only then were tested for boldness and handled. There was no habituation in the arena on the first day.

##### Open Field Test (OF)

Each rat was placed in an empty arena for 20 minutes. Each session was recorded, and the first 15 minutes were analyzed. Obtained movement patterns included total duration spent in the center, latency to first entering the center, frequency of entering the center, mean velocity, and total distance moved.

##### Boldness test

To examine whether individual baseline differences in boldness influence door­opening behavior, we tested the latency to approach the ledge of a halfway-opened cage as previously described by ^24^. Each cage was tested for 5 minutes for five days, and the timing of each rat to the ledge from the moment of opening the lid halfway was documented. In “ingroup” experiments, the boldness test was used to determine who would be the “free” and who would be the “trapped” rat in the HBT by calculating the total latencies of each rat. The rat with the lower total latency was classified as the “bold” rat and, therefore, was determined to be “free”. This test was not further analyzed and is not presented in this paper.

##### Helping Behavior Test (HBT)

In contrast with the previous protocol of the "separated" HBT, where rats were first trained to open the door in the non-separated HBT ^24^, here we overcame this training by allowing habituation to the restrainer during the habituation in the arena stages. This allowed the rats to be naive and without prior knowledge of door-opening when starting the experiment. The “separated” HBT was performed similarly to the original "nonseparated" HBT as described previously ^24^: 60-minute sessions were repeated daily over 12 days. During each session, a free rat was placed in an open arena containing a rat trapped inside a restrainer (25 by 8.75 by 7.5 cm, mechanical workshop for research and development, School of Chemistry, Tel-Aviv University). All free and trapped rats were age- and weight-matched. If the free rat had not opened the restrainer door during the 40 minutes since the beginning of the session, the investigator opened the restrainer door halfway, to a 45° angle; this was typically followed by the exit of the trapped rat and was aimed at preventing learned helplessness. Door-opening was counted as such when performed by the free rat before the halfway opening point. Rats that learned to open the restrainer and consistently opened it on the final three days of testing were labeled as "openers". In the “separated” HBT presented in this study, a plexiglass wall divided the arena into two compartments, thus preventing social contact following the release of the trapped rat. Two social conditions were tested: ingroup and outgroup. In the ingroup condition, the two rats (free and trapped) are from the same strain and have lived together in the same home cage for at least two weeks before the experiment. In the outgroup condition, the two rats are from different strains and were introduced to each other for the first time in the first session of the HBT. The free rat was introduced to the same stranger rat in the restrainer throughout the experiment.

#### Video Tracking

Sessions were recorded with a CCD color camera (Sony, China) connected to a video card (Geovision, Taipei, Taiwan) linked to a PC. Movement data were analyzed using Ethovision XT 15 video tracking software (Noldus Information Technology, Inc, Leesburg, VA). 8 male and 8 female LE rats tested in the ingroup condition, as well as 8 female LE rats tested in the outgroup condition were omitted from analysis of cumulative duration around the restrainer (Fig. 2E) due to a technical error.

#### Immunofluorescence

c-Fos staining procedures were described in detail in (Ben-Ami Bartal et al., 2021). At the end of the HBT, rats underwent one additional final session with a latched restrainer and were sacrificed within 60 minutes from the beginning of the session, at the peak expression of the early immediate gene product c-Fos. Rats were transcardially perfused with 1XPBS and freshly made 4% paraformaldehyde (PFA). Brains were then kept in 4% PFA overnight, sunk in 30% sucrose for additional 48 hours or until sunk, and stored at -80°C. They were later sliced at 40um (Leica CM3050 S; Leica 819 Low Profile Microtome Blades) and stored in cryoprotectant (Sucrose, Polyvinyl-pyrrolidone (PVP-40), 0.1M PB, Ethylene glycol) until stained for c-Fos. Free-floating sections were washed with 0.1M tris-buffered saline (TBS, 3X5’), incubated for 1 hour in 3% normal donkey serum (NDS) in 0.3% Triton X-100 in TBS (tTBS), and then transferred to rabbit anti-c-Fos (ab190289, Abcam, 1:1000; 1% NDS; 0.3% tTBS) in 4°C overnight. Sections were then washed in 0.1 M TBS (3X5’) and incubated for 2 hours in Alexa Fluor 488-conjugated donkey anti­rabbit (A-21206, 1:1000; 1% NDS; 0.3% tTBS). Sections were briefly washed in 0.1M TBS again (3X5’) and further stained in DAPI (1:40,000) for 10 min, then washed for an additional 15 min (3X5’). Lastly, all slides were mounted and coverslipped with 2.5% PVA/DABCO, dried overnight, and stored at 4°C until imaged. Immunostained tissue was imaged at 10* using a wide-field fluorescence microscope (Olympus ix83) and was processed (registered to brain atlas and quantified) in our open-source software Brainways ^22^.

#### Multiplex RNAscope

Brains from LE rats tested in the “separated” HBT in each social condition described above were used for fluorescent in situ hybridization (ISH) essays (ingroup, outgroup, and naive baseline, n = 6 each, half males and half females in each group. One rat was in the baseline condition was an outlier and thus excluded, resulting in n = 5 in this group). In order to reduce variability associated with door-opening in this relatively small sample, “openers” were selected for the ingroup and “non-openers” for the outgroup. As above, in the final session, restrainers were latched so that neural activity reflects an hour in the presence of a trapped rat. Rats were transcardially perfused with 1XPBS after the last HBT session. The brains were then snapfrozen and stored at -80°C until cryosectioning at 18//m. Coronal slices were collected on Superfrost Plus slides (Thermo Scientific) from each brain (4.20mm, 1.56mm, 0.00mm, -2.76mm, -3.36mm, -4.44mm, and -5.64mm from bregma) and underwent processing for single molecule fluorescent in situ hybridization (smFISH) assay. Slides were pre-treated with RNAscope Protease III reagent. smFISH was performed on slides using the RNAscope LS Multiplex Reagent Kit (Advanced Cell Diagnostics) and LS 4-Plex Ancillary Kit and Multiplex Reagent Kit on a robotic staining system (Leica BONDIII). RNAscope probes were Fos (ACD, 403598), OXTr (ACD, 483678), DrD2 (ACD, 315648), DrD1a (ACD, 317038). Images were acquired on a Vectra Polaris Automated Quantitative Pathology Imaging System (Akoya Biosciences) at 20* magnification. Cellular and subcellular detection was conducted using QuPath software and implemented into Brainways for image registration and brain-wide quantification.

#### Chemogenetic manipulations

##### Stereotaxic surgeries

Male LE rats (age 2-4 months) underwent stereotaxic injection of a viral vector containing an inhibitory (AAV8-hSyn-hM4D(Gi)-mCherry, n = 16; Addgene #50475) or an excitatory (AAV-hSynhM3D(Gq)-mCherry, n = 8; Addgene #50474) Designer Receptors Exclusively Activated by Designer Drugs (DREADDs) to the NAc (coordinates: A/P 1.5, M/L ±1.5, D/V -8.0). Rats were anesthetized with isoflurane (induction: 3-5%, maintenance: 1-3%) and mounted onto a stereotaxic frame. The skull was exposed following subcutaneous (s.c.) injection of Lidocaine (5mg/kg, 2%), and two small holes were made above the determined stereotaxic coordinates. A Hamilton Syringe with a needle of 33g containing the virus was used to inject 0.8­9 ul in each hemisphere. Following surgery, the rats were given s.c. injections of pain relievers (1mg/kg of Meloxicam 0.5%, 0.05mg/kg of Buprenorphine 0.3mg/ml) and Saline (10ml/kg) to ensure hydration. The rats were allowed two weeks to recover and then started the habituation and the HBT protocol, allowing three weeks in total for viral expression.

#### Behavioral testing

##### HBT with DREADDs manipulation

The DREADDs’ HBT habituation and testing were similar to the HBT procedure described above. In addition to the regular handling, rats were handled for intraperitoneal (i.p.) injection restraining (with a needle-less syringe). During testing of the first two groups (inhibition n = 8, excitation n = 8), a rat that performed three consecutive days of opening behavior was i.p. injected with Saline 0.9% (3 ml/kg) on the next day to exclude the alternative explanation that stress from the injection will alter the behavior. If opening continued, 3mg/kg Clozapine-N-Oxide (CNO; dissolved in 15ul DMSO 0.5% and filled with saline to injection volume; Cas No: 34233-69-7, Tocris) was i.p. administrated from the next day and every day for at least three days. This dose of CNO was selected as it was shown to be the highest effective and safe from side effects caused by CNO’s reverse-metabolism to clozapine ^57,58^. After all “openers” were administered with CNO for at least 3 days, all “non-opener” rats were treated with CNO for three additional days. Then, all rats were administered with saline for two final days. All injections were made 40 minutes before the testing session. The third group (inhibition n = 8) received CNO injections on days 1, 3, and 5 of the HBT (independently of opening behavior), with the same dosage as previous groups 40 minutes before the start of the session. On the rest of the days, the rats weren’t given any injections.

##### Control sessions

In order to investigate whether the CNO had a non-specific effect (on non-social/social-related behaviors), several control tests were conducted repeatedly over three days: with saline injection on the first day, CNO injection on the second day, and saline injection again on the third day. The control session included 5 minutes in an empty arena, 5 minutes with four pieces of apple (0.53 cm each) placed in the center of the arena, 5 minutes of interaction with the trapped rat, 5 minutes in an empty arena, 7 minutes with an empty restrainer, and 7 minutes with four pieces of apple placed in a restrainer.

##### Virus injection validation

After testing in the HBT and control sessions, rats underwent one final day of HBT with a latched restrainer as described in the former HBT procedure, after which the rats were sacrificed, and brains were obtained after perfusions with 1XPBS and 4% paraformaldehyde. Brains were cryosections in 40um and stained for c-Fos (rabbit anti-c-Fos, ab190289, Abcam; Alexa Fluor 488-conjugated donkey anti-rabbit, A-21206, Thermo Fischer) and DAPI as described in the “Immunofluorescence” section above. Co-labeling of c-Fos and mCherry was manually quantified in a representative area of 250um2 from the NAc from each rat.

#### Network analysis

Network graphs were generated by first obtaining a correlation matrix of c-Fos activity between all brain regions (using pairwise Pearson correlation coefficients). All significant correlations (p<0.05) were presented in a graphic form using Cytoscape ^59^. Correlation with significance values higher than the cutoff were set to one and the corresponding brain regions greater than 1 were considered connected to the network. Network parameters (clustering co-efficient, density) were analyzed with Cytoscape. Modularity was analyzed via Graph-tool.

#### Statistical analysis

Statistical details can be found in the Results section. For the RNAscope data, a threshold of 2 standard deviations was used to exclude outliers within each region, and regions that had less than 3 observations in one of the social conditions were also excluded. Missing data was then interpolated, and task PLS analysis was conducted as described before in ^26^. MATLAB code for running the Task PLS analysis is available for download from the McIntosh lab website. All means are reported as mean±SEM. Statistical analyses were performed using IBM SPSS Statistics version 28.0.1.0 (142), Graphpad Prism 9, and Python.

