## Supplementary Material for "Brain-wide activity-identity mapping of neural networks associated with prosocial motivation in rats"

### Supplementary information

Supplementary figures

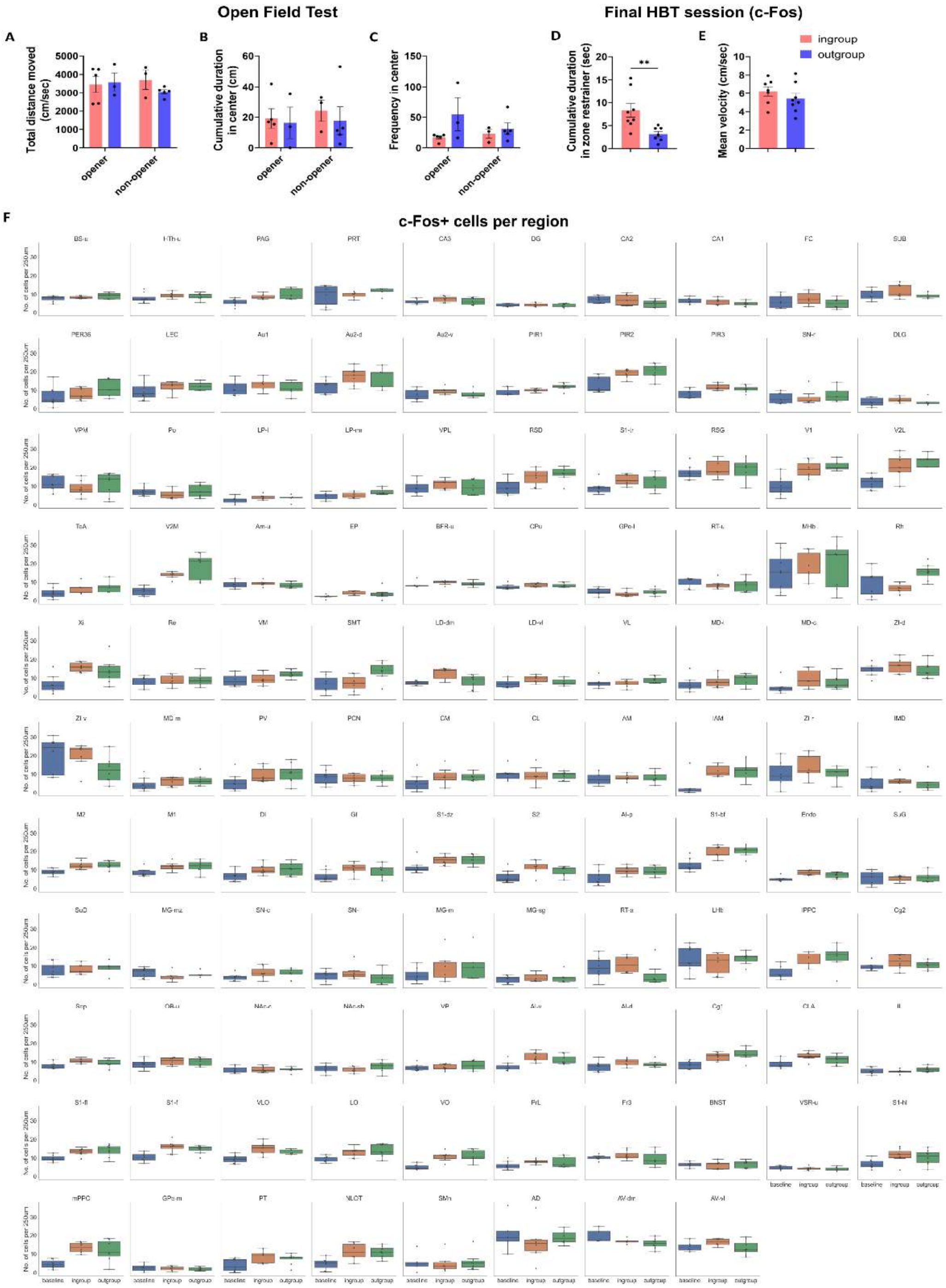


**Figure S1.** Supplementary behavioral and brain data from male SD rats in the "separated" paradigm. (**A-C**) Open Field Test results. (**D-E**) Movement data from the final HBT session, after which brains were obtained for further analysis. (**F**) Brain-wide c-Fos quantification via immunofluorescence staining per region and per social condition.

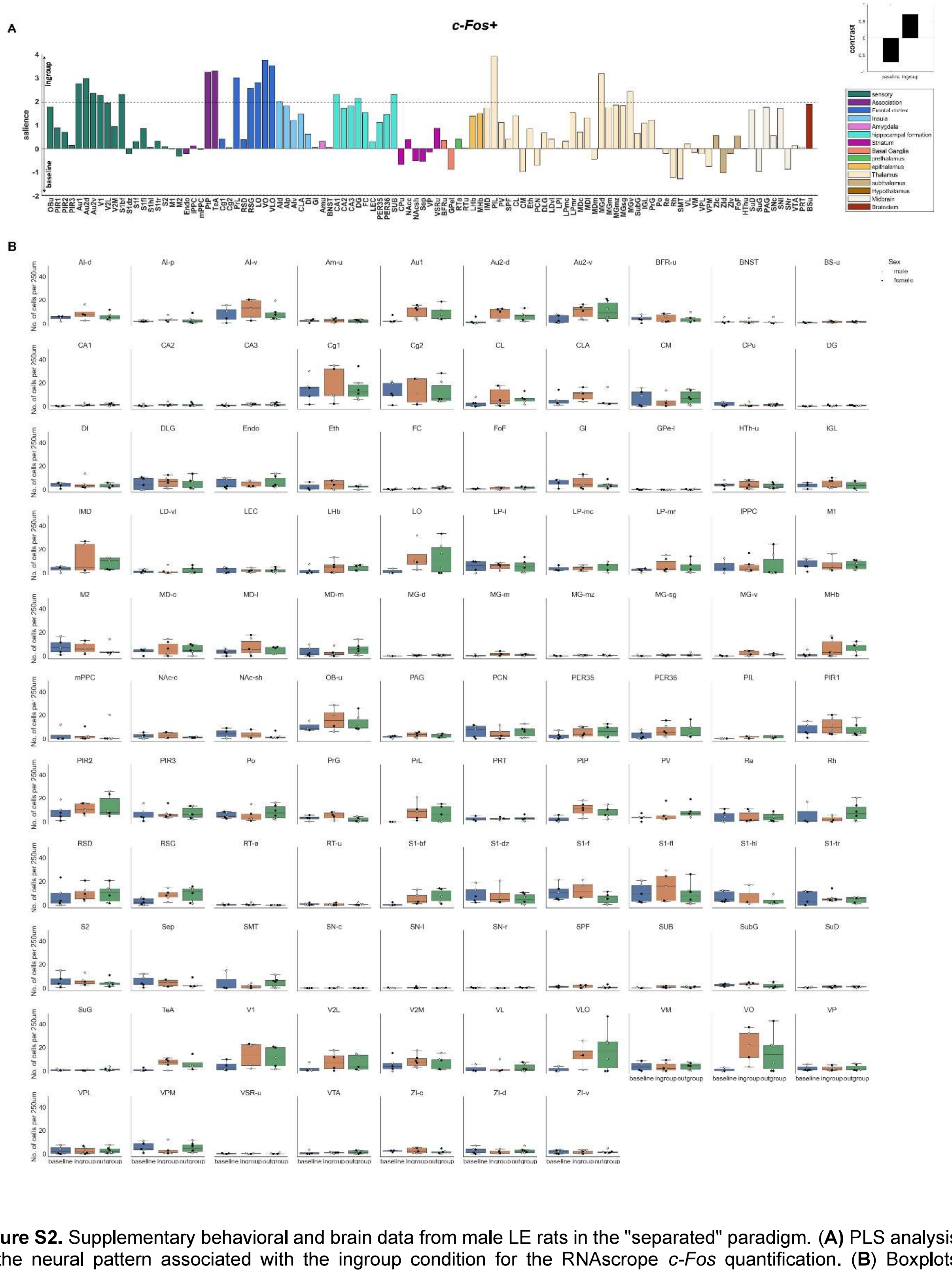


**Figure S2.** Supplementary behavioral and brain data from male LE rats in the "separated" paradigm. (**A**) PLS analysis of the neural pattern associated with the ingroup condition for the RNAscope *c-Fos* quantification. (**B**) Boxplots displaying RNAscope quantification of *c-Fos* per condition across brain regions.

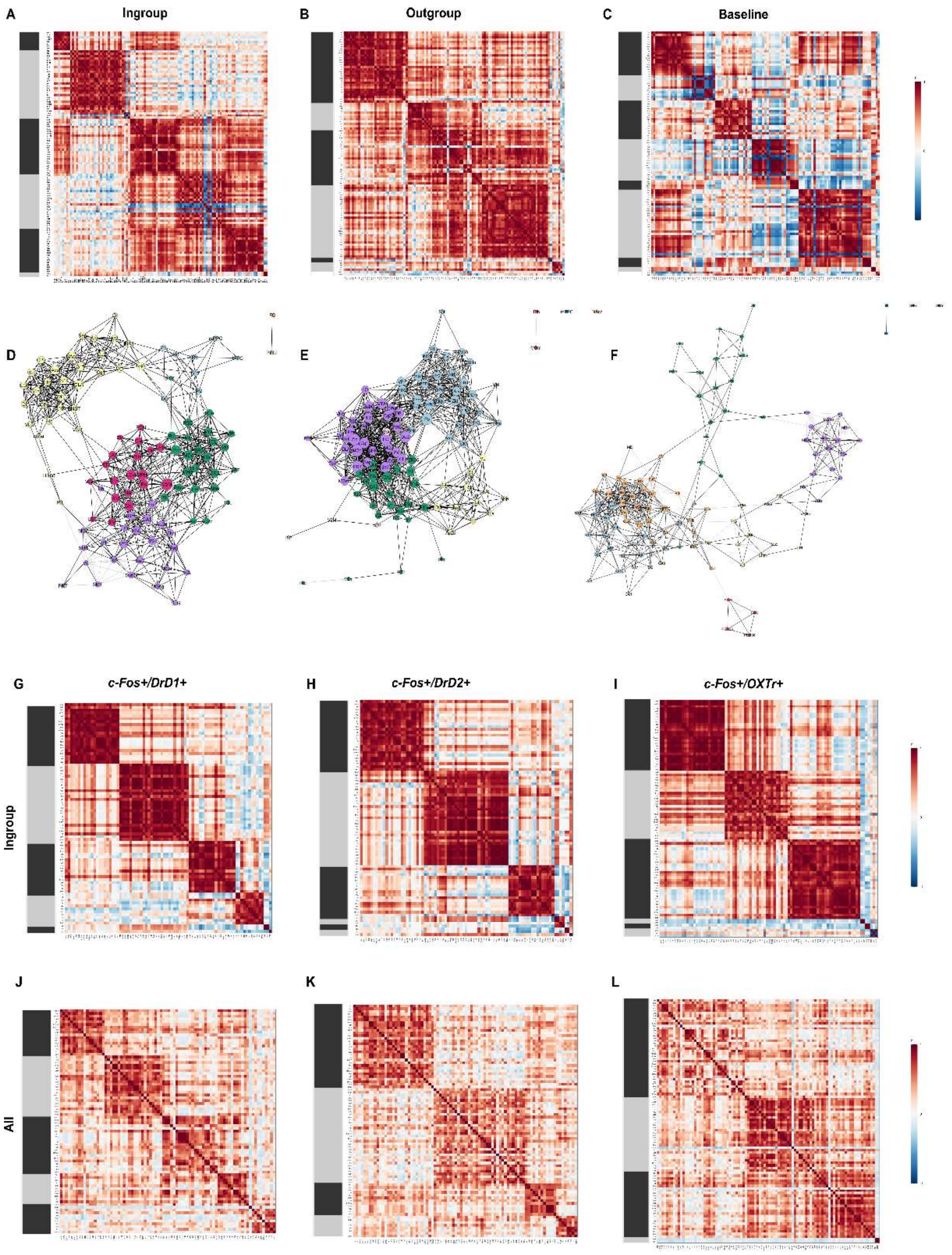


**Figure S3.** Neural networks associated with prosocial motivation. (**A-C**) Heatmaps generated from Pearson's correlations matrices across all brain regions in rats tested in the ingroup, outgroup and baseline conditions of the HBT (respectively), based on *c-Fos* RNAscope quantification. (**D-F**) Network graphs depicting the significant ($p<0.05$) inter-region correlations for the HBT ingroup, outgroup, and baseline conditions (respectively). Positive correlations are shown in black lines, negative correlations in gray lines. Circle color represents clusters identified via a Louvain algorithm, circle size represents the number of degrees for each region. (**G-I**) Heatmaps generated from Pearson's correlations matrices across all brain regions in rats tested the ingroup condition and (**J-L**) in all conditions of the HBT (ingroup, outgroup and baseline combined), based on (**G,J**) *c-Fos*$^{+}$*/Drd1*$^{+}$, (**H,K**) *c-Fos*$^{+}$*/Drd2*$^{+}$, and (**I,L**) *c-Fos*$^{+}$*/OXTr*$^{+}$ RNAscope quantification. Gray and black bars on the left of the matrices represent clusters identified via a Louvain algorithm.

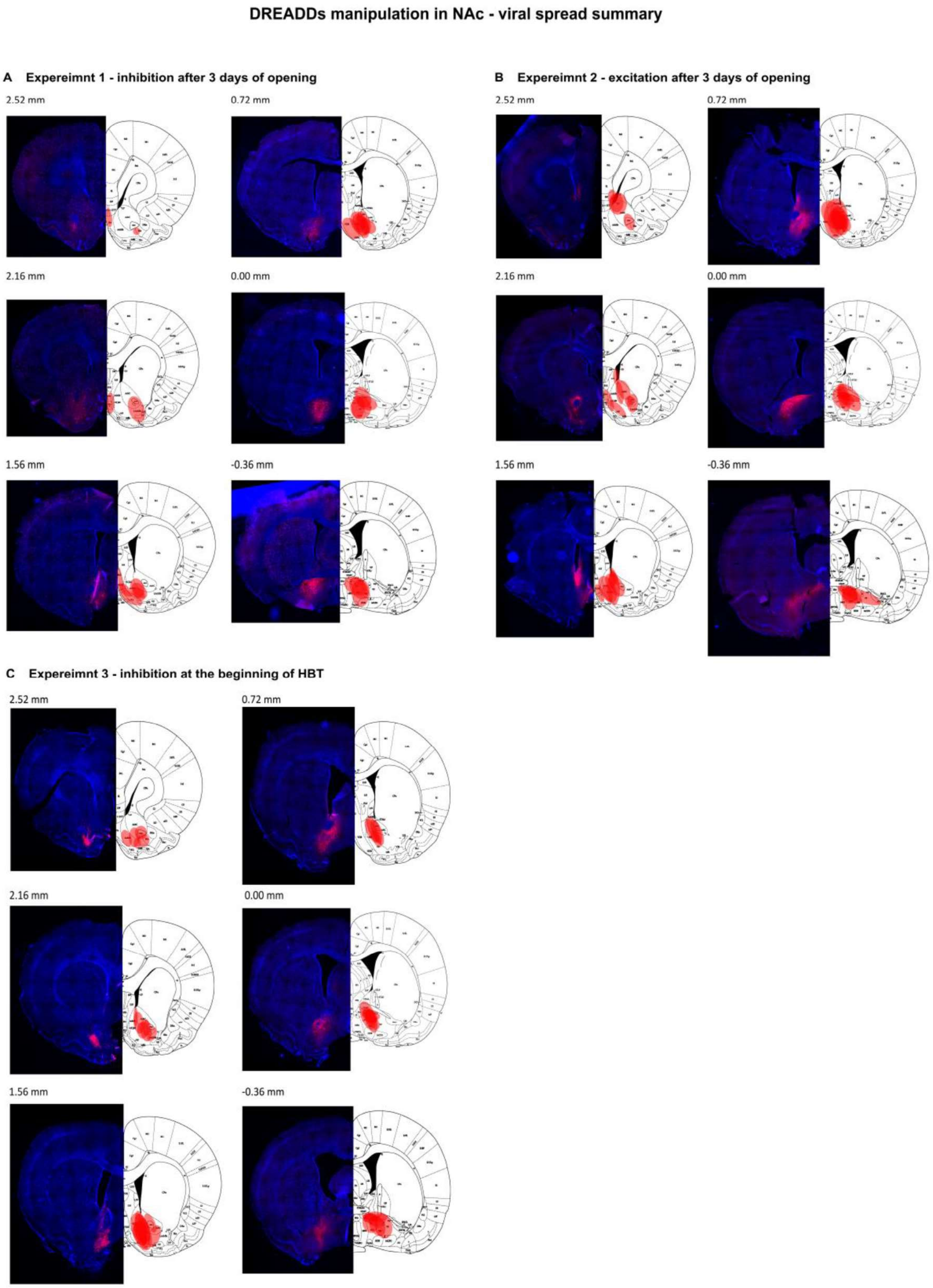


**Figure S4.** Viral spread in NAc chemogentic experiments. (**A**) Experiment 1: NAc inhibition after opening has been learned. (**B**) Experiment 2: NAc excitation after opening has been learned. (**C**) Experiment 3: Inhibition at the beginning of HBT.

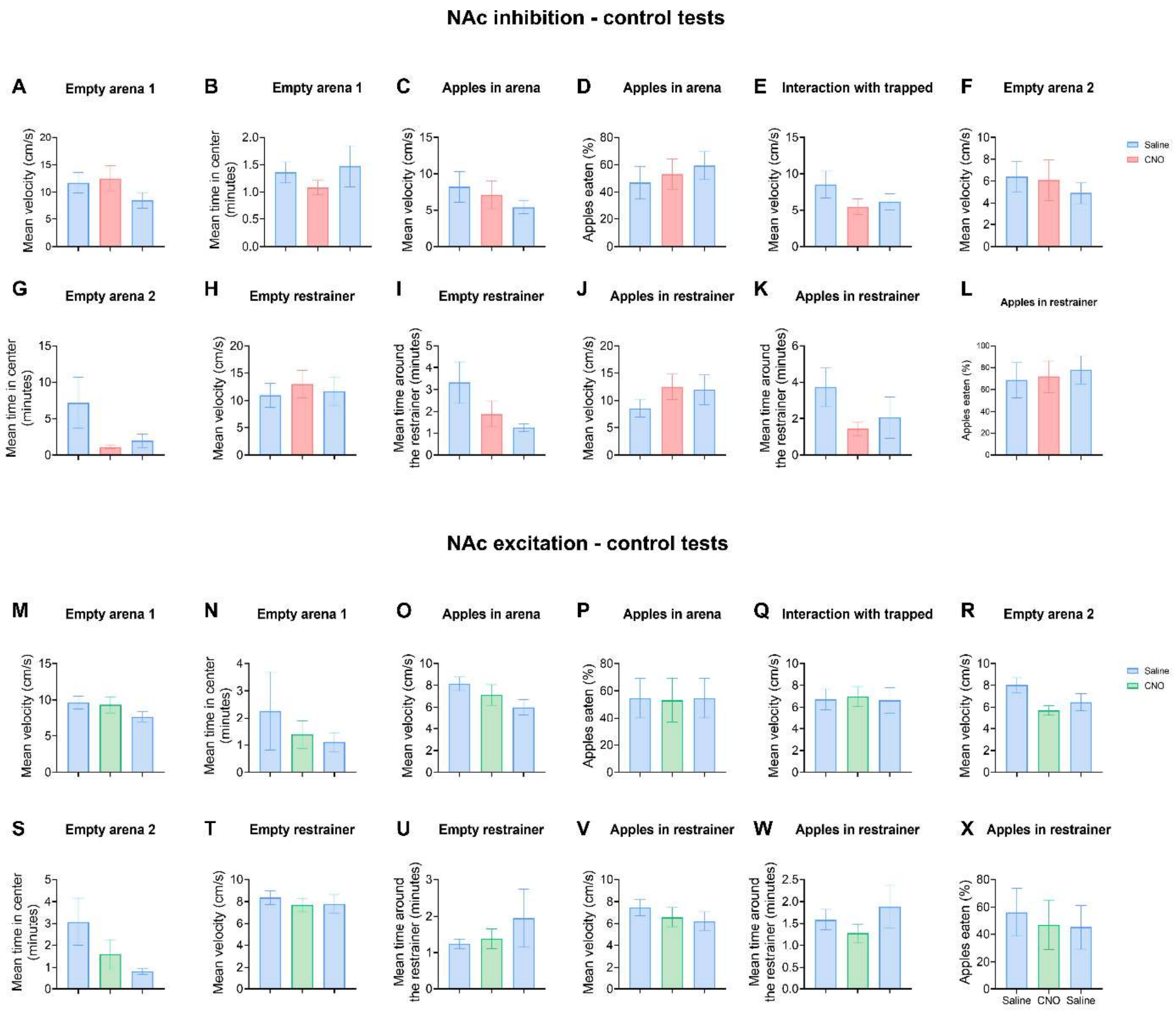


**Figure S5.** Control tests of NAc CNO-induced chemogenetic manipulation. Appearance of results is according to the testing order. CNO administration did not significantly affect locomotor activity and anxiety (inhibition **A-B, F-G**, excitation **M-N, R-S**), nonsocial reward-seeking or consumption (inhibition **C-D, J-L**, excitation **O-P, V-X**), interest in the restrainer (inhibition **H-I**, excitation **T-U**), or general social interaction (inhibition **E**, excitation **Q**).

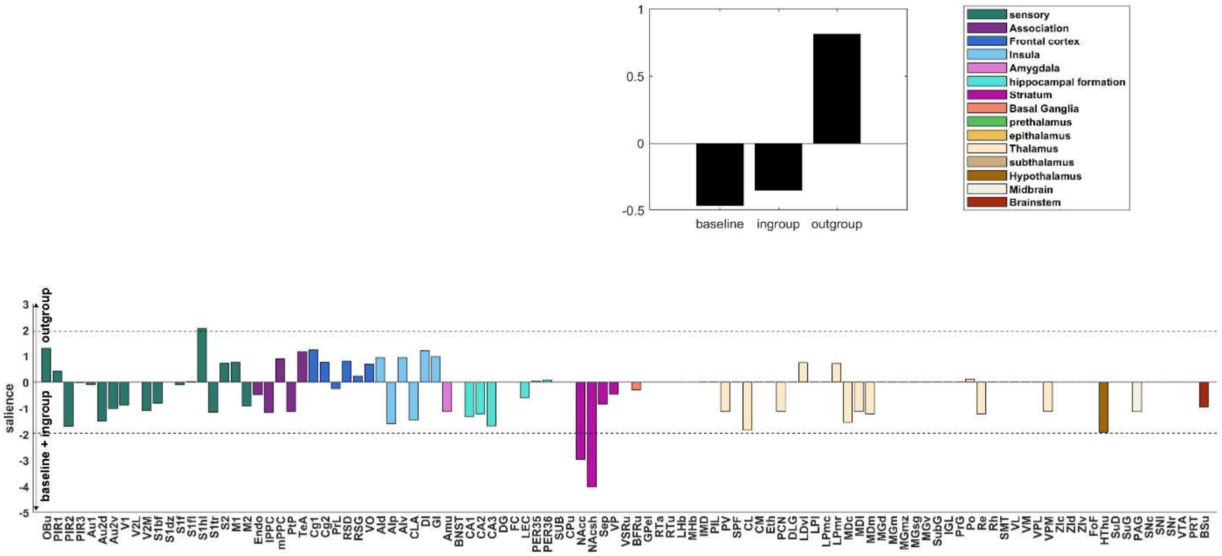


**Figure S6.** PLS analysis of % of cFos+*OXTr*+*DrD1*+ cells out of cFos+ cells.

### Supplementary tables

| acronym | name | acronym | name | acronym | name |
|---|---|---|---|---|---|
| AD | Anterodorsal thalamic nucleus | LPmc | Lateral posterior thalamic nucleus, mediocaudal part | S1dz | Primary somatosensory area, dysgranular zone |
| AId | Agranular insular cortex dorsal area | LPmr | Lateral posterior thalamic nucleus, mediorostral part | S1f | Primary somatosensory area, face representation |
| AIp | Agranular insular cortex, posterior area | lPPC | Parietal association cortex, lateral area | S1fl | Primary somatosensory area, forelimb representation |
| AIv | Agranular insular cortex, ventral area | M1 | Primary motor area | S1hl | Primary somatosensory area, hindlimb representation |
| AM | Anteromedial thalamic nucleus | M2 | Secondary motor area | S1tr | Primary somatosensory area, trunk representation |
| Amu | Amygdaloid area, unspecified | MDc | Mediodorsal thalamic nucleus, central part | S2 | Secondary somatosensory area |
| Ang | Angular thalamic nucleus | MDl | Mediodorsal thalamic nucleus, lateral part | Sag | Nucleus sagulum |
| Au1 | Primary auditory area | MDm | Mediodorsal thalamic nucleus, medial part | Sep | Septal region |
| Au2d | Secondary auditory area, dorsal part | MEC | Medial entorhinal cortex | SMn | Nucleus of the stria medullaris |
| Au2v | Secondary auditory area, ventral part | MGd | Medial geniculate body, dorsal division | SMT | Submedius thalamic nucleus |
| AVdm | Anteroventral thalamic nucleus, dorsomedial part | MGm | Medial geniculate body, medial division | SNc | Substantia nigra, compact part |
| AVvl | Anteroventral thalamic nucleus, ventrolateral part | MGmz | Medial geniculate body, marginal zone | SNl | Substantia nigra, lateral part |
| BFRu | Basal forebrain region, unspecified | MGsg | Medial geniculate body, suprageniculate nucleus | SNr | Substantia nigra, reticular part |
| BNST | Bed nucleus of the stria terminalis | MGv | Medial geniculate body, ventral division | SPF | Subparafascicular nucleus |
| BSu | Brainstem, unspecified | MHb | Medial habenular nucleus | STh | Subthalamic nucleus |
| CA1 | Cornu ammonis 1 | MO | Medial orbital area | SUB | Subiculum |
| CA2 | Cornu ammonis 2 | mPPC | Parietal association cortex, medial area | SubG | Subgeniculate nucleus |
| CA3 | Cornu ammonis 3 | NAcc | Nucleus accumbens, core | SuD | Deeper layers of the superior colliculus |
| Cg1 | Cingulate area 1 | NAcsh | Nucleus accumbens, shell | SuG | Superficial gray layer of the superior colliculus |
| Cg2 | Cingulate area 2 | NLOT | Nucleus of the lateral olfactory tract | TeA | Temporal association cortex |
| CL | Central lateral thalamic nucleus | OBu | Olfactory bulb, unspecified | V1 | Primary visual area |
| CLA | Claustrum | PAG | Periaqueductal gray | V2L | Secondary visual area, lateral part |
| CM | Central medial thalamic nucleus | PaS | Parasubiculum | V2M | Secondary visual area, medial part |
| CNIC | Inferior colliculus, central nucleus | PCN | Paracentral thalamic nucleus | VA | Ventral anterior thalamic nucleus |
| CPu | Caudate putamen | PER35 | Perirhinal area 35 | VL | Ventrolateral thalamic nucleus |
| DG | Dentate gyrus | PER36 | Perirhinal area 36 | VLO | Ventrolateral orbital area |
| DI | Dysgranular insular cortex | PF | Parafascicular thalamic nucleus | VM | Ventromedial thalamic nucleus |
| DLG | Dorsal lateral geniculate nucleus | PIL | Posterior intralaminar nucleus | VO | Ventral orbital area |
| DLO | Dorsolateral orbital area | PIR1 | Piriform cortex, layer 1 | VP | Ventral pallidum |
| ECIC | Inferior colliculus, external cortex | PIR2 | Piriform cortex, layer 2 | VPL | Ventral posterolateral thalamic nucleus |
| Endo | Endopiriform nucleus | PIR3 | Piriform cortex, layer 3 | VPM | Ventral posteromedial thalamic nucleus |
| EP | Entopeduncular nucleus | Pn | Pontine nuclei | VPpc | Ventral posterior nucleus of the thalamus, parvicellular part |
| Eth | Ethmoid-Limitans nucleus | Po | Posterior thalamic nucleus | VSRu | Ventral striatal region, unspecified |
| FC | Fasciola cinereum | Pot | Posterior thalamic nuclear group, triangular part | VTA | Ventral tegmental area |
| FoF | Fields of Forel | PP | Peripeduncular nucleus | Xi | Xiphoid thalamic nucleus |
| Fr3 | Frontal association area 3 | PrG | Pregeniculate nucleus | ZIA11 | Zona incerta, A11 dopamine cells |
| GI | Granular insular cortex | PrL | Prelimbic area | ZIA13 | Zona incerta, A13 dopamine cells |
| GPel | Globus pallidus external, lateral part | PrS | Presubiculum | ZIc | Zona incerta, caudal part |
| GPem | Globus pallidus external, medial part | PRT | Pretectal region | ZId | Zona incerta, dorsal part |
| HThu | Hypothalamic region, unspecified | RT | Reticular (pre)thalamic nucleus | ZIr | Zona incerta, rostral part |
| IAM | Interanteromedial thalamic nucleus | PT | Parataenial thalamic nucleus | ZIv | Zona incerta, ventral part |
| IGL | Intergeniculate leaflet | PtP | Parietal association cortex, posterior area | | |
| IL | Infralimbic area | PV | Paraventricular thalamic nuclei (anterior and posterior) | | |
| IMD | Intermediodorsal thalamic nucleus | Re | Reuniens thalamic nucleus | | |
| IP | Interpeduncular nucleus | Rh | Rhomboid thalamic nucleus | | |
| LDdm | Laterodorsal thalamic nucleus, dorsomedial part | RRe | Retroreuniens thalamic nucleus | | |
| LDvl | Laterodorsal thalamic nucleus, ventrolateral part | RSD | Retrosplenial dysgranular area | | |
| LEC | Lateral entorhinal cortex | RSG | Retrosplenial granular area | | |
| LHb | Lateral habenular nucleus | RTa | Reticular (pre)thalamic nucleus, auditory segment | | |
| LO | Lateral orbital area | RTu | Reticular (pre)thalamic nucleus, unspecified | | |
| LPl | Lateral posterior thalamic nucleus, lateral part | S1bf | Primary somatosensory area, barrel field | | |

**Table S1. Brain regions' abbreviations.**

| Region | *c-Fos+/DrD1+* | | *c-Fos+/DrD2+* | | *c-Fos+/OXTr+* | |
|---|---|---|---|---|---|---|
| | Average | SD | Average | SD | Average | SD |
| **amygdala** | **1.89** | **2.54** | **3.65** | **4.92** | **6.39** | **5.89** |
| Amu | 1.96 | 2.02 | 1.46 | 1.53 | 8.23 | 7.29 |
| BNST | 1.82 | 3.05 | 5.83 | 8.30 | 4.54 | 4.49 |
| **association** | **1.46** | **2.05** | **1.03** | **1.01** | **3.27** | **5.40** |
| Endo | 4.47 | 5.51 | 3.83 | 3.04 | 5.57 | 3.95 |
| lPPC | 1.38 | 2.55 | 0.27 | 0.49 | 1.98 | 4.83 |
| mPPC | 1.19 | 1.65 | 0.54 | 0.78 | 4.21 | 9.92 |
| PtP | 0.17 | 0.34 | 0.27 | 0.32 | 2.72 | 5.84 |
| TeA | 0.09 | 0.18 | 0.26 | 0.43 | 1.87 | 2.48 |
| **brainstem** | **0.13** | **0.21** | **4.13** | **3.26** | **2.17** | **1.98** |
| BSu | 0.13 | 0.21 | 4.13 | 3.26 | 2.17 | 1.98 |
| **epithalamus** | **0.78** | **1.32** | **0.50** | **0.87** | **2.76** | **5.38** |
| LHb | 1.37 | 2.28 | 0.47 | 0.73 | 3.80 | 8.02 |
| MHb | 0.18 | 0.36 | 0.52 | 1.00 | 1.71 | 2.73 |
| **frontal** | **1.11** | **1.67** | **1.45** | **1.31** | **2.81** | **3.32** |
| Cg1 | 1.64 | 2.55 | 1.64 | 0.99 | 3.61 | 3.35 |
| Cg2 | 1.51 | 2.11 | 1.88 | 1.32 | 1.78 | 1.62 |
| LO | - | - | 1.18 | 1.22 | 1.35 | 2.23 |
| M1 | 0.57 | 0.57 | 0.88 | 0.75 | 3.56 | 3.18 |
| M2 | 1.03 | 1.54 | 1.16 | 0.92 | 6.22 | 5.97 |
| PrL | 1.83 | 3.31 | 2.44 | 3.04 | - | - |
| RSD | 0.51 | 0.76 | 1.21 | 1.08 | 1.87 | 3.49 |
| RSG | 0.69 | 0.85 | 1.18 | 1.15 | 1.71 | 3.50 |
| VO | - | - | - | - | 2.41 | 3.24 |
| **hippocampus** | **0.30** | **0.56** | **0.48** | **0.67** | **4.97** | **5.82** |
| CA1 | 0.37 | 0.65 | 0.65 | 0.83 | 5.31 | 4.64 |
| CA2 | 0.25 | 0.44 | 0.16 | 0.55 | 3.46 | 5.37 |
| CA3 | 0.26 | 0.25 | 0.31 | 0.41 | 5.79 | 8.02 |
| DG | 0.39 | 0.68 | 0.46 | 0.83 | 3.13 | 2.85 |
| FC | 0.11 | 0.32 | 0.24 | 0.54 | 8.00 | 11.32 |
| LEC | 0.32 | 0.49 | 0.83 | 0.93 | 4.82 | 5.53 |
| PER35 | 0.35 | 0.70 | 0.40 | 0.51 | 5.44 | 5.60 |
| PER36 | 0.51 | 0.99 | 0.42 | 0.55 | 3.72 | 4.33 |
| SUB | 0.12 | 0.49 | 0.83 | 0.90 | 5.08 | 4.73 |
| **hypothalamus** | **0.55** | **0.64** | **5.94** | **4.64** | **4.72** | **3.60** |
| HThu | 0.55 | 0.64 | 5.94 | 4.64 | 4.72 | 3.60 |
| **insula** | **1.02** | **1.77** | **2.20** | **2.08** | **4.06** | **4.64** |
| AId | 0.59 | 1.05 | 3.29 | 2.96 | 4.99 | 6.64 |
| AIp | 0.56 | 0.89 | 0.74 | 0.82 | 5.01 | 6.64 |
| AIv | 0.64 | 1.23 | 2.49 | 2.83 | 3.86 | 3.64 |
| CLA | 1.63 | 2.74 | 3.01 | 2.61 | 0.61 | 1.01 |
| DI | 1.02 | 1.89 | 2.05 | 1.96 | 5.39 | 5.83 |
| GI | 1.66 | 2.84 | 1.59 | 1.31 | 4.48 | 4.06 |
| **midbrain** | **0.05** | **0.10** | **3.21** | **4.55** | **1.15** | **1.48** |
| PAG | 0.16 | 0.36 | 2.86 | 1.83 | 5.19 | 5.88 |
| PRT | 0.23 | 0.45 | 0.97 | 0.80 | 0.80 | 1.23 |
| SNc | 0.00 | 0.00 | 9.15 | 15.95 | 0.00 | 0.00 |
| SNl | 0.00 | 0.00 | 0.00 | 0.00 | 0.00 | 0.00 |
| SNr | 0.00 | 0.00 | 2.99 | 5.48 | 0.00 | 0.00 |
| SuD | 0.00 | 0.00 | 2.49 | 3.53 | 1.70 | 2.73 |
| SuG | 0.00 | 0.00 | 0.63 | 1.27 | 0.00 | 0.00 |
| VTA | 0.00 | 0.00 | 6.56 | 7.52 | 1.53 | 2.02 |
| **sensory** | **1.02** | **1.30** | **0.53** | **0.55** | **3.07** | **3.06** |
| Au1 | 0.48 | 0.59 | 0.45 | 0.44 | 1.80 | 2.75 |
| Au2d | 0.20 | 0.26 | 0.39 | 0.47 | 2.35 | 3.67 |
| Au2v | 0.34 | 0.49 | 0.54 | 0.53 | 3.17 | 4.07 |
| OBu | 3.04 | 4.13 | 0.72 | 0.92 | 13.19 | 5.24 |
| PIR1 | 2.72 | 3.57 | 0.29 | 0.32 | 2.19 | 2.24 |
| PIR2 | 3.08 | 4.19 | 0.54 | 0.59 | 4.27 | 4.14 |
| PIR3 | 3.02 | 3.64 | 1.26 | 0.91 | 5.04 | 4.72 |

| Region | *c-Fos+/DrD1+* | | *c-Fos+/DrD2+* | | *c-Fos+/OXTr+* | |
|---|---|---|---|---|---|---|
| | Average | SD | Average | SD | Average | SD |
| S1bf | 0.53 | 0.57 | 0.39 | 0.40 | 2.60 | 2.73 |
| S1dz | 0.51 | 0.66 | 0.73 | 0.76 | 1.98 | 2.49 |
| S1f | 0.76 | 0.85 | 0.42 | 0.36 | 2.41 | 2.81 |
| S1fl | 0.37 | 0.48 | 0.55 | 0.35 | 2.48 | 2.17 |
| S1hl | 0.30 | 0.41 | 0.38 | 0.47 | 1.31 | 1.27 |
| S1tr | 0.60 | 0.64 | 0.51 | 0.61 | 1.87 | 3.09 |
| S2 | 0.54 | 0.54 | 0.54 | 0.48 | 2.97 | 3.29 |
| V1 | 0.16 | 0.19 | 0.50 | 0.45 | 1.94 | 3.49 |
| V2L | 0.06 | 0.12 | 0.18 | 0.30 | 1.53 | 2.19 |
| V2M | 0.56 | 0.82 | 0.67 | 0.94 | 1.02 | 1.62 |
| **striatum** | **12.44** | **9.53** | **21.18** | **9.82** | **4.57** | **3.23** |
| Sep | 1.91 | 2.38 | 7.54 | 4.55 | 7.94 | 3.68 |
| BFRu | 5.14 | 3.28 | 5.82 | 2.95 | 8.09 | 4.70 |
| CPu | 27.90 | 15.20 | 43.20 | 8.27 | 2.16 | 2.09 |
| GPel | 9.41 | 7.27 | 9.65 | 11.58 | 0.42 | 1.73 |
| NAcc | 19.79 | 13.85 | 43.13 | 11.98 | 1.31 | 1.95 |
| NAcsh | 20.22 | 17.70 | 20.34 | 10.97 | 6.67 | 4.46 |
| VP | 11.77 | 11.55 | 14.12 | 9.96 | 7.94 | 3.93 |
| VSRu | 3.38 | 5.00 | 25.65 | 18.31 | 2.02 | 3.30 |
| **subthalamus** | **0.14** | **0.35** | **14.12** | **12.74** | **2.04** | **3.28** |
| FoF | 0.00 | 0.00 | 37.79 | 33.89 | 0.68 | 1.92 |
| ZIc | - | - | 10.05 | 6.67 | 1.56 | 1.97 |
| ZId | 0.35 | 0.75 | 4.51 | 4.93 | 3.95 | 6.01 |
| ZIv | 0.07 | 0.29 | 4.13 | 5.48 | 1.97 | 3.22 |
| **thalamus** | **0.25** | **0.55** | **0.22** | **0.41** | **2.28** | **4.46** |
| CL | 0.28 | 0.60 | 0.06 | 0.19 | 6.61 | 11.47 |
| CM | 0.22 | 0.36 | 0.23 | 0.50 | 8.16 | 16.74 |
| DLG | 0.02 | 0.06 | 0.02 | 0.06 | 1.16 | 3.00 |
| Eth | 0.00 | 0.00 | 0.00 | 0.00 | 2.14 | 4.06 |
| IGL | 0.00 | 0.00 | 0.00 | 0.00 | 0.51 | 1.40 |
| IMD | 0.22 | 0.63 | 0.00 | 0.00 | 6.37 | 11.36 |
| LDvl | 0.60 | 1.27 | 0.34 | 1.04 | 2.70 | 7.04 |
| LPl | 0.00 | 0.00 | 0.04 | 0.13 | 1.09 | 2.08 |
| LPmc | 0.00 | 0.00 | 0.00 | 0.00 | 0.98 | 2.26 |
| LPmr | 0.43 | 0.93 | 1.99 | 3.67 | 1.96 | 4.33 |
| MDc | 0.72 | 1.67 | 0.00 | 0.00 | 4.66 | 11.29 |
| MDl | 0.74 | 1.86 | 0.03 | 0.13 | 3.99 | 8.95 |
| MDm | 0.13 | 0.38 | 0.07 | 0.28 | 3.44 | 6.11 |
| MGd | 0.00 | 0.00 | 0.00 | 0.00 | 0.00 | 0.00 |
| MGm | 0.00 | 0.00 | 0.00 | 0.00 | 0.33 | 0.94 |
| MGmz | 0.00 | 0.00 | 0.00 | 0.00 | 0.00 | 0.00 |
| MGsg | 0.00 | 0.00 | 0.00 | 0.00 | 0.00 | 0.00 |
| MGv | 0.00 | 0.00 | 0.00 | 0.00 | 0.00 | 0.00 |
| PCN | 0.32 | 0.64 | 0.00 | 0.00 | 3.28 | 7.63 |
| PIL | 0.00 | 0.00 | 0.00 | 0.00 | 0.00 | 0.00 |
| Po | 0.24 | 0.45 | 0.05 | 0.12 | 0.61 | 1.64 |
| PrG | 0.14 | 0.41 | 0.10 | 0.35 | 0.28 | 0.68 |
| PV | 0.15 | 0.51 | 0.23 | 0.42 | 11.76 | 14.05 |
| Re | 0.33 | 0.62 | 0.06 | 0.19 | 4.20 | 5.68 |
| Rh | 0.06 | 0.24 | 0.00 | 0.00 | 0.51 | 1.21 |
| RTa | 0.37 | 1.52 | 0.00 | 0.00 | 0.00 | 0.00 |
| RTu | 1.09 | 2.88 | 0.00 | 0.00 | 0.28 | 1.15 |
| SMT | 0.05 | 0.16 | 0.00 | 0.00 | 0.05 | 0.16 |
| SPF | 0.00 | 0.00 | 2.54 | 3.60 | 0.96 | 2.31 |
| SubG | 0.00 | 0.00 | 0.59 | 1.40 | 0.34 | 0.82 |
| VL | 0.58 | 1.20 | 0.01 | 0.03 | 4.94 | 11.82 |
| VM | 1.61 | 1.91 | 1.15 | 1.90 | 2.95 | 5.49 |
| VPL | 0.04 | 0.12 | 0.00 | 0.00 | 0.89 | 1.52 |
| VPM | 0.09 | 0.16 | 0.00 | 0.00 | 2.36 | 6.41 |

**Table S2. % of co-labeled cells out of total c-Fos+ cells per region**

**Supplementary Videos**

**Video S1. Example video of the "separated" HBT.** Door-opening by a female LE rat tested in the ingroup condition.

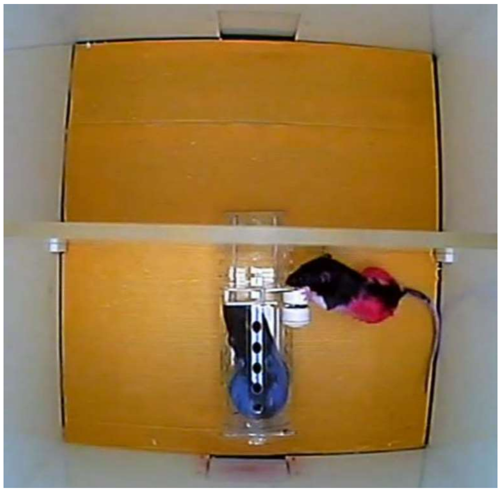
